# Neural population dynamics in dorsal premotor cortex underlying a reach decision

**DOI:** 10.1101/2022.06.30.497070

**Authors:** Pierre O Boucher, Tian Wang, Laura Carceroni, Gary Kane, Krishna V Shenoy, Chandramouli Chandrasekaran

## Abstract

We investigated if a dynamical systems approach could help understand the link between decision-related neural activity and decision-making behavior, a fundamentally unresolved problem. The dynamical systems approach posits that neural dynamics can be parameterized by a state equation that has different initial conditions and evolves in time by combining at each time step, recurrent dynamics and inputs. For decisions, the two key predictions of the dynamical systems approach are that 1) initial conditions substantially predict subsequent dynamics and behavior and 2) inputs should combine with initial conditions to lead to different choice-related dynamics. We tested these predictions by investigating neural population dynamics in the dorsal premotor cortex (PMd) of monkeys performing a red-green reaction time (RT) checkerboard discrimination task where we varied the sensory evidence (i.e., the inputs). Prestimulus neural state, a proxy for the initial condition, predicted poststimulus neural trajectories and showed organized covariation with RT. Furthermore, faster RTs were associated with faster pre- and poststimulus dynamics as compared to slower RTs, with these effects observed within a stimulus difficulty. Poststimulus dynamics depended on both the sensory evidence and initial condition, with easier stimuli and “fast” initial conditions leading to the fastest choice-related dynamics whereas harder stimuli and “slow” initial conditions led to the slowest dynamics. Finally, changes in initial condition were related to the outcome of the previous trial, with slower pre- and poststimulus population dynamics and RTs on trials following an error as compared to trials following a correct response. Together these results suggest that decision-related activity in PMd is well described by a dynamical system where inputs combine with initial conditions that covary with eventual RT and previous outcome, to induce decision-related dynamics.

## 1. Introduction

There are 10 minutes to make it to the airport but the GPS says you’re still 12 minutes away. Seeing a yellow light in the distance you quickly floor it. You get to the intersection only to realize you have run a red light. The sight of the lights result in patterns of neural activity that respectively lead you to respond quickly to your environment (i.e., speed up when you see the yellow) and process feedback (i.e., slow down after running the red). This process of choosing, performing, and altering actions in response to sensory cues and context is termed perceptual decision-making (Cisek, 2012; Kiani et al., 2013; Brody and Hanks, 2016; Gold and Shadlen, 2007; Brunton et al., 2013).

Research in invertebrates (Briggman et al., 2005; Kato et al., 2015), rodents (Hanks et al., 2015; Guo et al., 2014), monkeys (Roitman and Shadlen, 2002; Churchland et al., 2008), and humans (Pereira et al., 2021; Kelly and O’Connell, 2013) has attempted to understand the neural basis for decision-making. Barring few exceptions (Okazawa et al., 2021; Mante et al., 2013; Thura et al., 2020), emphasis has been placed on understanding and characterizing single neuron responses in decision-related brain regions (Roitman and Shadlen, 2002; Churchland et al., 2008; Thura and Cisek, 2014; Chandrasekaran et al., 2017). However, how these processes manifest in neural population dynamics to mediate decision-making behavior, especially in reaction time (RT) tasks, is largely unclear. In this study, we address this gap by investigating if a “dynamical systems” approach, originally posited in motor planning studies, can provide a mechanistic understanding of decision-related dynamics and behavior (Churchland et al., 2006; Afshar et al., 2011; Shenoy et al., 2013).

The dynamical systems approach (Vyas et al., 2020a; Shenoy et al., 2013; Remington et al., 2018a) posits that neural population activity, *X* is governed by a state equation of the following form:

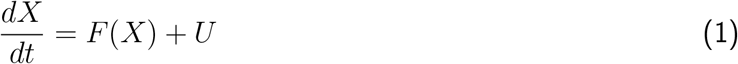

Where *F* is a function that represents the recurrent dynamics (i.e., synaptic input) in the region of interest, *U* is the input to the system from neurons outside the region of interest, and *X*_0_ is the initial condition for these dynamics. The function *F* is usually considered to be fixed for a given brain area in a task, and *U* is variable depending on various task contingencies (e.g., sensory evidence). In this framework, dynamics for every trial are dependent on both the initial conditions and inputs and this ultimately leads to distinct behavior on every trial.

The attractiveness of the dynamical systems approach is that it provides a powerful and simplified mechanistic basis for understanding the link between time-varying, heterogeneous activity of neural populations and behavior (Afshar et al., 2011; Kaufman et al., 2014; Elsayed et al., 2016). For example, in studies of motor planning, the position and velocity of the neural population dynamics relative to the mean trajectory at the time of the ‘go’ cue (i.e., initial condition or *X*_0_) explained considerable variability in RTs (Afshar et al. 2011, see Fig. 1A). Similarly, in studies of timing, the initial condition encoded the perceived time interval and predicted the speed of subsequent neural dynamics and the reproduced time interval (Remington et al. 2018b, see Fig. 1B). In the same study, an input depending on a task contingency (“gain”) also altered the speed of dynamics (Fig. 1B).

**Figure 1:**
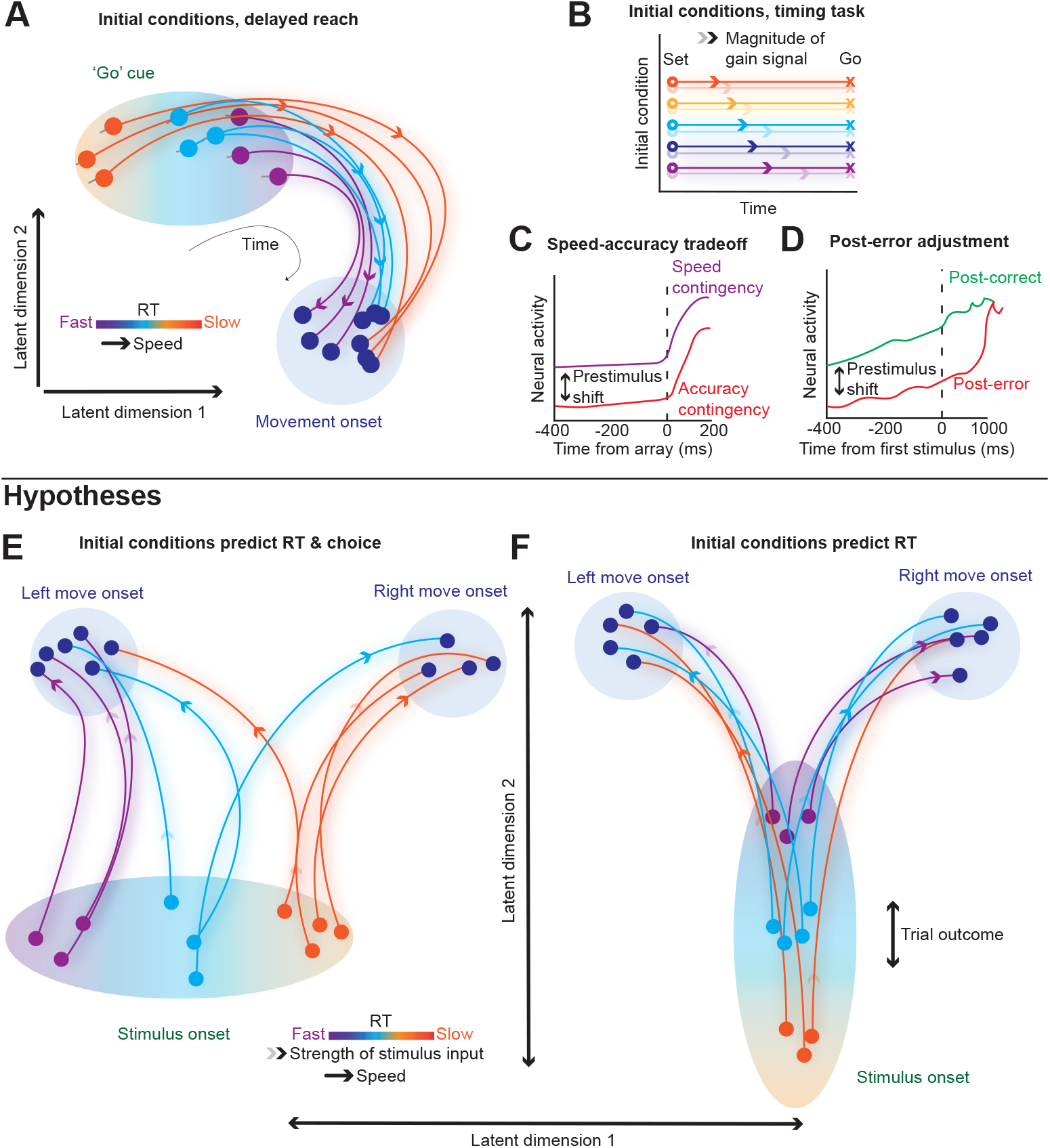
Initial conditions and inputs predict subsequent neural dynamics and behavior. (**A**) The initial condition hypothesis from delayed reach experiments (Afshar et al., 2011) posits that the position and velocity of a neural state at the time of the ‘go’ cue (“initial condition”) negatively correlates with RT. That is for faster RT trials, neural state at the time of the go cue is 1) further along (“position”) relative to the mean neural trajectory and thus closer to the movement initiation state and 2) has a greater rate of change in the direction of the mean neural trajectory (“velocity”). (**B**) The neural population state at the end of a perceived time interval and a gain modifier actuates the initial conditions (Set, circles) determining the speed (arrows) of subsequent dynamics and therefore when an action is produced (Go, X’s) (Remington et al., 2018b). (**C & D**) Prestimulus neural activity differs for speed and accuracy contingencies for speed-accuracy tradeoff tasks (Heitz and Schall, 2012) or after correct and error trials (Thura et al., 2017). (**E**) Biased initial conditions predict both *RT* and *choice* (*X*_0_ ~ *RT, choice*) and combine with sensory evidence to lead to decisions. Initial neural states vary trial-to-trial, and are closer to the movement onset state for one choice (here left). Trials with neural states closer to a left movement onset at stimulus onset will have faster RTs and RTs will be slower for right choices. Trial outcomes have no effect on initial conditions in this model as initial conditions largely reflect a reach bias. (**F**) Initial conditions solely predict RT (*X*_0_ ~ *RT*). The position of the initial condition before checkerboard onset is closer to a movement initiation state and the velocity of the dynamics are faster for fast RTs compared to slow RTs. Previous outcomes shift these initial conditions such that the dynamics are either faster or slower, leading to faster or slower RTs respectively. Overall dynamics depend on both the initial conditions and the sensory evidence. Current population state at stimulus onset/go cue (dots within an ellipse) evolves along trajectories of varying speed (color bars in **A & E**; apply to **A, B, E** and **F**) as set by the initial conditions (**A**) and also inputs after stimulus onset (**E & F**). In **E** and **F** light/dark opacity of the arrowhead indicates weak/strong stimulus input.

Our goal here was to expand on these findings from motor planning and timing studies and investigate if a dynamical systems approach with varying initial conditions and inputs could provide a mechanistic understanding of neural population activity underlying decisions. Support for varying initial conditions in decisions comes from speed-accuracy tradeoff (SAT) and post-error adjustment experiments. In these studies, prestimulus neural activity is different for fast vs. slow blocks (Murphy et al., 2016; Bogacz et al., 2010) and depends on the outcome of the previous trial, respectively (Purcell and Kiani, 2016; Thura et al., 2017), see Fig. 1C/D. Similarly, a large body of research emphasizes that the rate at which choice-selective activity emerges is dependent on the strength of the sensory evidence (Roitman and Shadlen, 2002; Chandrasekaran et al., 2017; Hanks et al., 2015; Coallier et al., 2015, e.g., auditory pulses, random dot motion, static red-green checkerboards, etc.). Thus, based on this prior work, we hypothesize that a dynamical system for decision-making should have the following properties: 1) the initial condition, as indexed by the prestimulus neural population state, predicts poststimulus decision-related neural dynamics and behavior and 2) the speed of choice-selective dynamics after stimulus onset should depend on the strength of the sensory evidence and the initial conditions. The initial neural state will either predict both RT-and choice-(Fig. 1E, *X*_0_ ~ *RT, choice*) or only RT-related poststimulus dynamics and behavior (Fig. 1F, *X*_0_ ~ *RT*), with the dynamics in the latter hypothesis dependent upon previous trial outcomes.

We tested these predictions by examining firing rates of neurons recorded in dorsal premotor cortex (PMd) of monkeys performing a red-green RT perceptual decision-making task (Chandrasekaran et al., 2017). First, analysis of the state space trajectories suggested that neural population dynamics were ordered pre- and poststimulus as a function of RT. Subsequent KiNeT analysis (Remington et al., 2018b) of the dynamics of these trajectories suggested that faster RTs were associated with faster pre- and poststimulus dynamics as compared to slower RTs and such effects were observed within a stimulus difficulty. Decoding and regression analyses further revealed that prestimulus neural state, that is the *initial condition*, only predicted RT but not the eventual choice consistent with the hypothesis shown in Fig. 1F. The speed of the poststimulus dynamics that led to the eventual choice **jointly depended on the initial condition and the sensory evidence**, with choice-related signals emerging faster for easier compared to harder trials but also modulated by the initial condition. Initial conditions and choice-related dynamics depended on the outcome of the previous trial with pre- and poststimulus dynamics slower on trials following an error as compared to trials following a correct response. Our results are a significant and important expansion of the observations of Afshar et al. (2011), that the prestimulus position and velocity of the neural trajectories in state space (i.e., initial conditions) are correlated with RT, as we demonstrate that 1) both inputs and initial conditions jointly control dynamics, and 2) that changes in the initial conditions are dependent upon previous outcomes. Together these results suggest that decision-related activity in PMd is captured by a dynamical system composed of initial conditions, that predict RT and are dependent upon previous outcome, and inputs (i.e., sensory evidence) which combine with initial conditions to induce choice-related dynamics.

## 2. Results

### 2.1. Decision-related behavior is dependent on sensory evidence and internal state

We trained two macaque monkeys (O and T) to discriminate the dominant color of a central, static checkerboard composed of red and green squares (Fig. 2A). Fig. 2B depicts the trial timeline. The trial began when the monkey held the center target and fixated on the fixation cross. After a short randomized holding time (300-485 ms), a red and a green target appeared on either side of the central hold (target configurations were randomized). After an additional randomized target viewing time drawn from a censored exponential distribution (400-1000 ms), the checkerboard appeared. The monkey’s task was to reach to, and touch the target corresponding to the dominant color of the checkerboard. While animals were performing the task, we measured the arm and eye movements of the monkeys. We identified RTs as the first time when hand speed exceeded 10% of maximum speed during a reach. If the monkey correctly performed a trial, he was rewarded with a drop of juice and a short inter-trial interval (ITI, 300 to 600 ms across sessions) whereas if he made an error it led to a longer timeout ITI (ranging from ~ 1500 ms to ~ 3500 ms). Using timeouts for errors encouraged animals to prioritize accuracy over speed.

**Figure 2:**
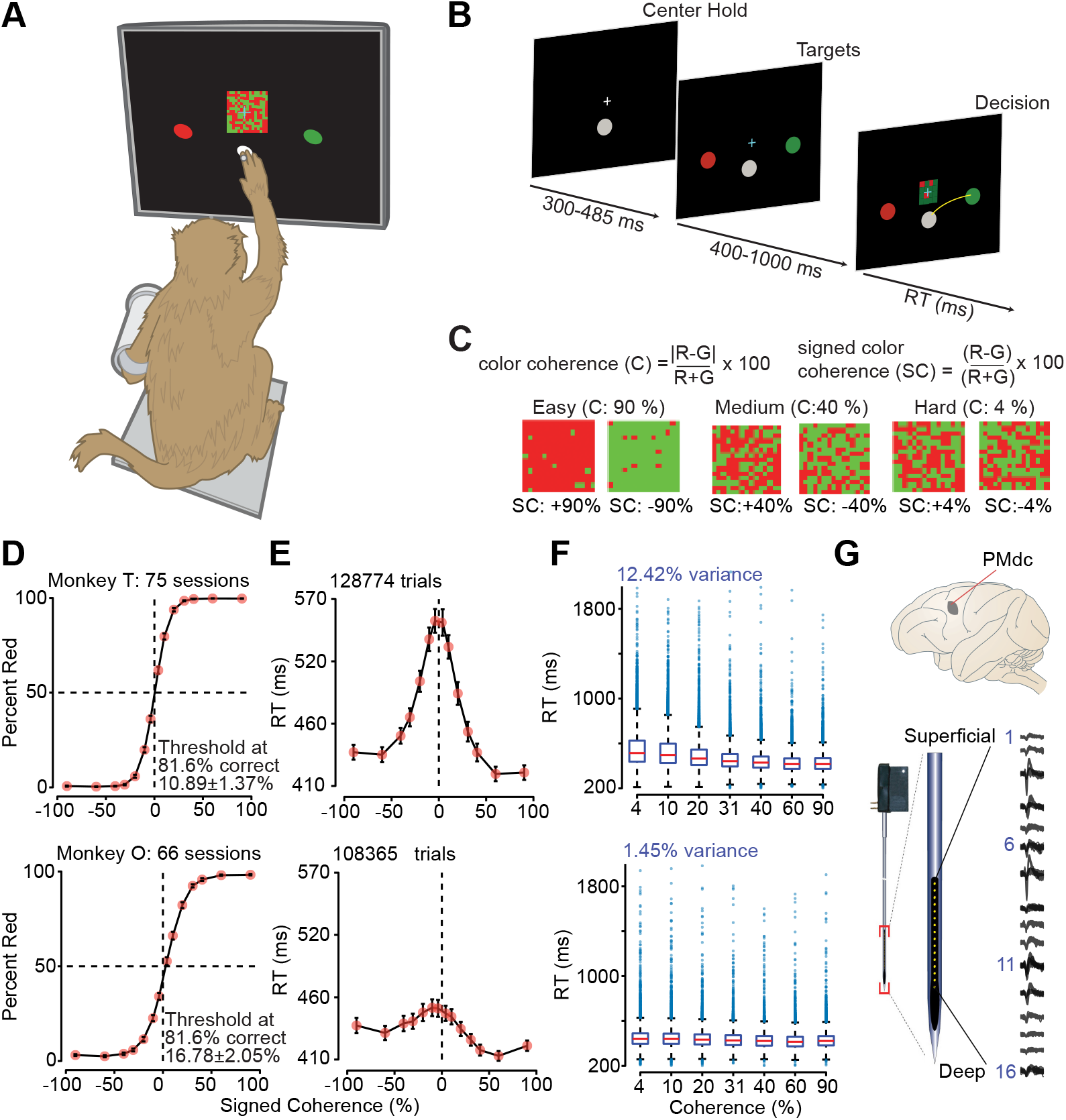
Monkeys can discriminate red-green checkerboards and demonstrate rich variability in RTs between and within stimulus coherences. (**A**) An illustration of the setup for the behavioral task. We loosely restrained the arm the monkey was not using with a plastic tube and cloth sling. A reflective infrared bead was taped on the middle digit of the active hand to be tracked in 3D space. We used the measured hand position to mimic a touch screen and to provide an estimate of instantaneous arm position; eye position was tracked using an infrared reflective mirror placed in front of the monkey’s nose. (**B**) Timeline of the discrimination task. (**C**) Examples of different stimuli used in the experiment parameterized by the color coherence of the checkerboard cue. Positive values of signed coherence (*SC*) denote more red (R) than green (G) squares and vice versa. (**D**) Psychometric curves, percent responded red, and (**E**) RTs (correct and incorrect trials) as a function of the percent *SC* of the checkerboard cue, over sessions of the two monkeys (T: 75 sessions; O: 66 sessions). Dark orange markers show measured data points along with 2 × SEM estimated over sessions (error bars lie within the marker for many data points). The black line segments are drawn in between these measured data points to guide the eye. Discrimination thresholds measured as the color coherence level at which the monkey made 81.6% correct choices are also indicated. Thresholds were estimated using a fit based on the cumulative Weibull distribution function. (**F**) Box-and-whisker plot of RT as a function of unsigned checkerboard coherence with outliers plotted as blue circles. Note large RT variability within and across coherences. (**G**) The recording location, caudal PMd (PMdc), indicated on a macaque brain, adapted from Ghazanfar and Santos (2004). Single and multi-units in PMdc were primarily recorded by a 16 electrode (150-µm interelectrode spacing) U-probe (Plexon, Inc., Dallas, TX, United States); example recording depicted.

We used 14 levels of sensory evidence referred to as signed color coherence (‘*SC*’, Fig. 2C) as it’s *dependent* on the actual dominant color of the checkerboard. Unsigned coherence (‘*C*’, Fig. 2C), which refers to the strength of stimuli, is *independent* of the actual dominant color of the checkerboard. Thus, there are 7 levels of *C*.

The behavioral performance of the monkeys depended on the signed coherence. In general, across all sessions, monkeys made more errors when discriminating stimuli with near equal combinations of red and green squares (Fig. 2D). We fit the proportion correct as a function of unsigned coherence using a Weibull distribution function to estimate slopes and psychometric thresholds (average *R*^2^; T: 0.99 (over 75 sessions); O: 0.98 (over 66 sessions); Threshold - Mean *±* SD: T: 10.89 *±* 1.37%, O: 16.78 *±* 2.05% slope (*β*, Mean *±* SD over sessions, T: 1.26 *±* 0.18, O: 1.10 ± 0.14).

As expected, monkeys were generally slower for more ambiguous checkerboards (Fig. 2E). However, per monkey regressions using unsigned coherence (*log*_10_(*C*)) to predict RTs only explained ~ 12.4% and ~ 1.5% of RT variability, for monkeys T and O respectively. These results suggest that while there is RT variability induced by differences in the stimulus evidence, there is also an internal source of RT variability. Indeed, as the box plots in Fig. 2F show, a key feature of the monkeys’ behavior is that RTs are quite variable within a coherence, even for the easiest ones. In the subsequent sections, we investigated if a dynamical system parameterized by initial conditions and inputs could explain RT variability and choice behavior.

### 2.2. Single unit prestimulus firing rates covary with RT and poststimulus activity is input dependent

Our database for understanding the neural population dynamics underlying decision-making consists of 996 units (546 units in T and 450 units in O, including both single neurons and multi-units, 801 single neurons) recorded from PMd of the two monkeys over 141 sessions. We chose units as they were well isolated from other units/separated from noise and modulated activity in at least one task epoch. A unit was categorized as a single neuron by a combination of spike sorting and if inter-spike-interval violations were minimal (≤ 1.5% of inter-spike-intervals were ≤ 1.5 ms; median across single neurons: 0.28%).

Fig. 3 shows the smoothed (30 ms Gaussian) firing rates of five example units recorded in PMd aligned to checkerboard onset and organized either by coherence and choice, plotted until the median RT (Fig. 3A), or organized by RT and choice, plotted until the midpoint of the RT bin (Fig. 3B). Many units showed classical ramp-like firing rates (Shadlen and Newsome 1996, 2001; Roitman and Shadlen 2002; Hanks et al. 2014; Latimer et al. 2015, see Fig. 3, top three rows). However, many neurons demonstrated complex, time-varying patterns of activity that included increases and decreases in firing rate that covaried with stimulus difficulty, choice and RT (Fig. 3, bottom 2 rows) (Meister et al., 2013; Mante et al., 2013; Jun et al., 2010). Additionally, each of the, albeit curated, neurons in Fig. 3B demonstrated prestimulus firing rate covariation with RT implying variable initial conditions that ultimately factor into RTs.

**Figure 3:**
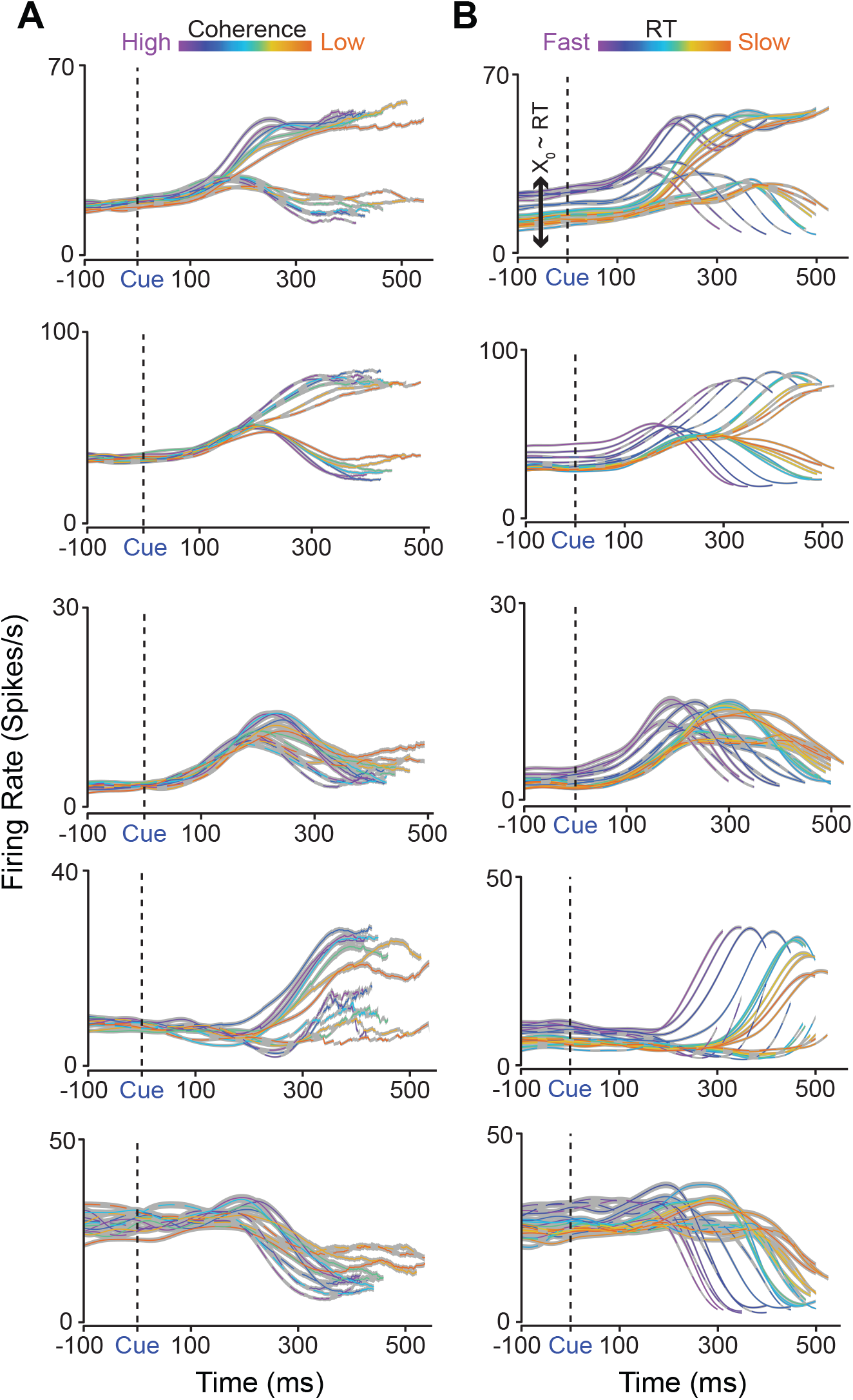
Firing rates of a heterogeneous population of PMd neurons are modulated by the input (i.e., strength of the sensory evidence) and the initial conditions (prestimulus firing rate) covaries with RT before stimulus onset. (**A, B**) Firing rate activity across (**A**) 7 levels of color coherence and (**B**) 11 RT bins and both action choices (right - dashed, left - solid) of 5 example units in PMd from monkeys T and O aligned to stimulus onset (Cue/vertical dashed black line). Firing rates are plotted until the median RT of each color coherence and until the midpoint of each RT bin (notice slightly different lengths of lines). Color bars indicate the level of difficulty for the coherence (violet - mostly one color, orange - nearly even split of red and green squares) or RT speed. Gray shading is *SEM*. In **A**, the firing rates separate faster for easier choices compared to harder choices, and in **B**, the same neurons show prestimulus modulation as a function of RT (*X*_0_ ~ *RT*).

Thus, these example units provide preliminary support for our hypothesis that variable initial conditions, combined with sensory evidence as input can explain decision-related dynamics and behavior. In the next sections, we used dimensionality reduction, decoding, and regression analyses to further interrogate how RT and choice were represented in the shared, time-varying, and heterogeneous activity of these neurons.

### 2.3. Principal component analysis reveals prestimulus population state covariation with RT

The single unit examples shown in Fig. 3 support the proposition that the initial conditions, or population neural dynamics just before stimulus onset, should strongly account for RT variability and this effect should be observed within a stimulus difficulty. To visualize if this was the case, we performed a principal component analysis (PCA) on trial-averaged firing rate activity (again smoothed with a 30 ms Gaussian) windowed about checkerboard onset, organized by overlapping RT bins, 11 levels representing a spectrum from faster to slower RTs (300-400 ms, 325-425 ms, …, to 600-1000 ms), and both reach directions (Fig. 4A, B). For this analysis, we pooled all trials (including both correct and wrong trials) across all the different stimulus coherences and sorted by RT and choice before averaging.

**Figure 4:**
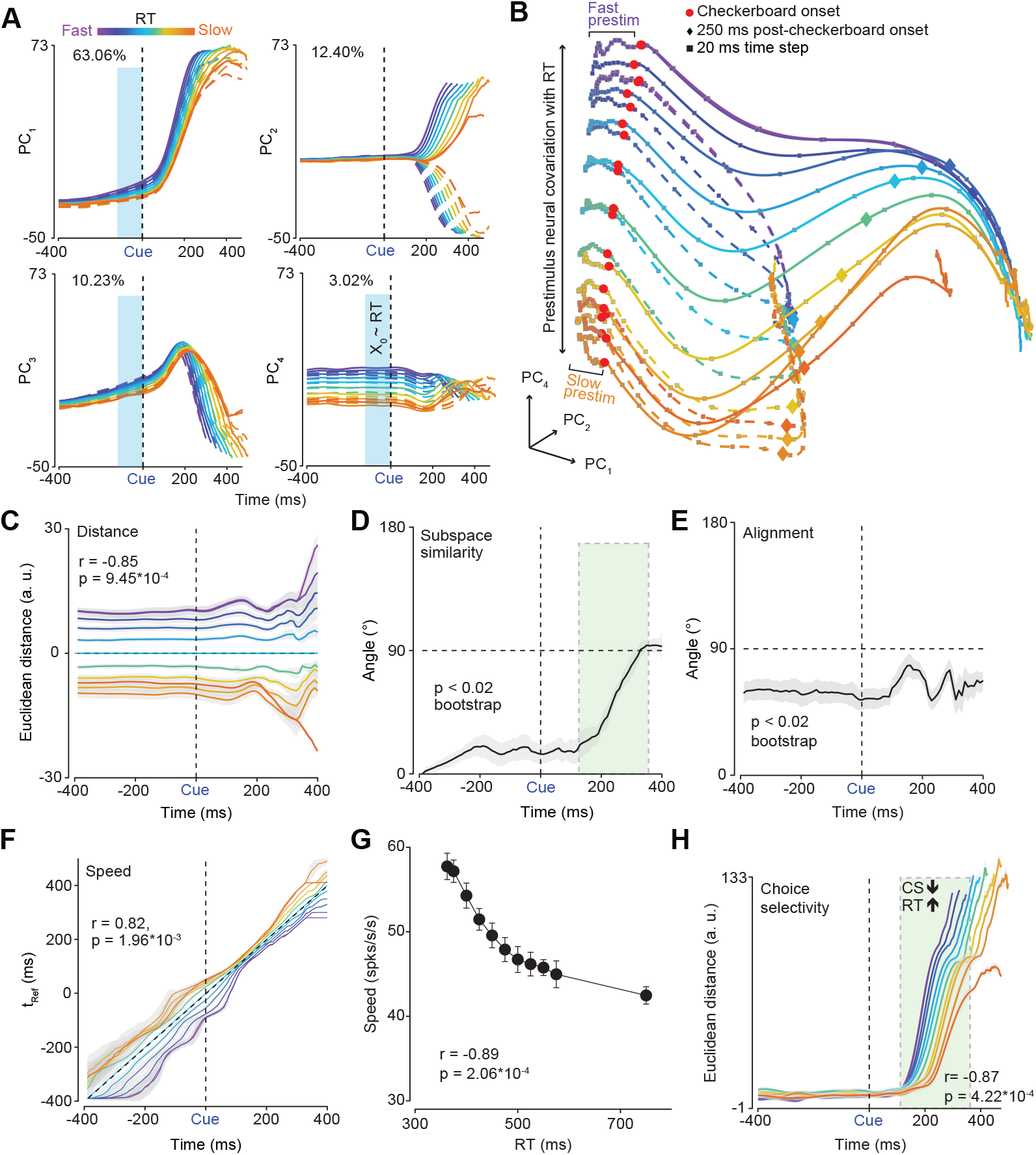
Prestimulus population firing rates covary with RT. (**A**) The first four PCs (*PC*_1,2,3,4_) of trial averaged firing rates organized across 11 RT bins (violet - fastest bin to orange - slowest bin, both reach directions (right - dashed lines, left - solid lines), and aligned to checkerboard onset. Percent variance explained by each PC is indicated at the top of each plot. (**B**) State space trajectories of the 1st, 2nd and 4th PCs (*PC*_1,2,4_) aligned to checkerboard onset (red dots). Prestimulus neural activity robustly separates as a function of RT bin. Diamonds and squares, color matched to their respective trajectories, indicate 250 ms post-checkerboard onset and 20 ms time steps respectively. Note that faster RT trajectories appear to move faster in the prestimulus period than slower RTs (“fast/slow prestim”, also see **G**). (**C**) “KiNeT distance” analysis showing that trajectories are consistently spatially organized before and after stimulus onset and correlated with RT. (**D**) Angle between subspace vector at each timepoint and subspace vector at the first timepoint (- 400ms). The angle between subspace vectors is largely consistent but the space rotates as choice signals emerge (green highlight box). (**E**) Average relative angle between adjacent trajectories for each timepoint. The angles between adjacent trajectories were largely less than 90^*°*^ for the prestimulus period but approach orthogonality as choice signals emerge poststimulus. (**F**) “KiNeT Time to reference” (t_Ref_, relative time at which a trajectory reaches the closest point in Euclidean space to the reference trajectory) analysis shows that trajectories for faster RTs reach similar points on the reference trajectory (cyan, middle trajectory) earlier than trajectories for slower RTs. This result suggests that the dynamics for faster RTs are closer to a movement initiation state than slower RTs. (**G**) Average scalar speed for the prestimulus period (−400 to 0 ms epoch) as a function of RT bin. Firing rates across the population change faster (both increases and decreases) for faster RTs and slower for slower RTs. Error bars are bootstrap *SEM*. (**H**) Choice selectivity signal measured as the Euclidean distance in the first six dimensions between the two reach directions for each RT bin aligned to checkerboard onset. The rate at which Choice selectivity (CS) emerges is faster for faster RTs compared to slower RTs (green highlight box). In **C & F** the x-axis is labelled “Time (ms)”, this should be understood as time on the reference trajectory. *Abbreviations:* Checkerboard onset - Cue & vertical black dashed line, a. u. - Arbitrary units.

To identify the number of relevant dimensions for describing this data, we used a principled approach developed in Machens et al. (2010) (see 4.10 for details). Firing rates on every trial in PMd during this task can be thought of as consisting of a combination of signal (i.e., various task related variables) and noise contributions from sources outside the task such as spiking noise, for example. Trial averaging reduces this noise but nevertheless when PCA is performed it returns a principal component (PC) space that captures variance in firing rates due to the signal and variance due to residual noise (“signal+noise PCA”). Ideally, we only want to assess the contributions of the signal to the PCA, but this is not possible for trial-averaged or non-simultaneously recorded data. To circumvent this issue and determine the number of signal associated dimensions, the method developed in Machens et al. (2010) estimates the noise contributions by performing a PCA on the difference between single trial estimates of firing rates, to obtain a “noise” PCA. Components from the signal+noise PCA and the noise PCA were compared such that only signal+noise dimensions that were significantly greater than the noise dimensions (identified as the first point where the signal+noise variance was significantly lower than the noise variance by at least 3 × SEM) were included in further analyses. The assumption here is that the dimensions above the noise are largely dominated by the signal and the dimensions below the noise are largely noise dimensions. This analysis yielded six PCs that explained *>* 90% of the variance in firing rates (Fig. S1).

Fig. 4A plots the first four PCs obtained from this PCA. What is apparent in Fig. 4A is that the prestimulus state strongly covaries with RT but not with choice. In particular, barring component 2, which seems to be most strongly associated with choice, PCs 1, 3, and 4 showed covariation between the prestimulus state and RT (Fig. 4A, highlighted with light blue rectangles) — consistent with the rich covariation between RT and prestimulus firing rates in the single neuron examples shown in Fig. 3B. Visualizing PCs 1, 2, and 4 in a state space plot further supported this observation (Fig. 4B). In this state space plot, both position and the velocity of the prestimulus state appear to covary with RT. For instance, prestimulus trajectories for the fastest RTs are 1) spatially separated, and 2) appear to have covered more distance along the paths for movement initiation by the time of checkerboard onset than the prestimulus trajectories for the slowest RTs (small squares which denote 20 ms time steps are more spread out for faster versus slower trajectories, Fig. 4B). In contrast, only a modest separation by choice occurs before stimulus onset.

Note, such covariation between prestimulus neural state and RT was not an artifact of pooling across all the different stimulus difficulties and was observed even within a level of stimulus coherence (note similarities between Fig. 4B & Fig. 6C). We discuss this further in section 2.6 where we analyzed the joint effects of inputs and initial conditions.

Collectively, the visualization using PCA firmly suggests that prestimulus state predicts poststimulus dynamics and covaries with RT but not the eventual choice. In subsequent sections, we used various analyses to further understand if these data could be understood through the lens of a dynamical system that has varying initial conditions, and inputs. We first examined how initial conditions control the dynamics of decision-making, and then how they combined with inputs to drive decisions.

### 2.4. Position and ‘velocity’ of initial condition correlate with poststimulus dynamics and RT

The dynamical systems perspective predicts that poststimulus dynamics and behavior depend upon the position and velocity of prestimulus neural trajectories in state space (i.e., initial conditions) (Shenoy et al., 2013; Vyas et al., 2020a). Position is the instantaneous location in a high-dimensional state space of neural activity (i.e., firing rate of neurons) and velocity is a directional measure of how fast these positions are changing over time (i.e., directional rate of change from one neural state to the next). We used the Kinematic analysis of Neural Trajectories (KiNeT) approach recently developed by Remington et al. (2018b) to test this prediction. KiNeT measures the spatial ordering of trajectories and how each trajectory evolves in time, all with respect to a reference trajectory. Please refer to Fig. S4 for a visualization of how KiNeT analyses are performed and see 4.11 for a full description of KiNeT calculations.

First, we used KiNeT to assess if position of the initial conditions was related to RT. If the position of the initial condition covaries with RT then we would expect a lawful ordering of neural trajectories organized by RT bin, otherwise they would lie one on top of the other indicating a lack of spatial organization. Thus, we examined the spatial ordering of six-dimensional neural trajectories grouped by RT bins for each reach direction. We estimated the signed minimum Euclidean distance at each point for the trajectory relative to a reference trajectory (the middle RT bin, cyan, for that reach direction, Fig. 4C). Trajectories were 1) organized by RT with trajectories for faster and slower RT bins on opposite sides of the the reference trajectory, and 2) the relative ordering of the Euclidean distance with respect to the reference trajectory was also lawfully related to RT (Fig. 4C) as measured by a correlation between RT and the signed Euclidean distance at −100 ms before checkerboard onset (r = −0.85, p = 9.45 × 10^−4^). These data are consistent with the prediction that the position of the initial condition correlates with RT.

Second, we examined if the relative ordering of trajectories by RT in the prestimulus period predicted the ordering of poststimulus trajectories by measuring the “subspace similarity” angle, and average “alignment”. Subspace similarity is a measure of how six-dimensional neural trajectories rotate as a subspace in time in relation to the first timepoint. Alignment measures the degree to which neural trajectories diverge from one another in state space by estimating the average angle of the normalized vectors between pairwise adjacent trajectories at each timepoint. The null hypothesis is that prestimulus dynamics do not have consistent spatial ordering and adjacent trajectories are rapidly changing which would lead to changes in the subspace angle between adjacent time points and large changes in the alignment of adjacent trajectories. Alternatively, if prestimulus dynamics predict ordering of poststimulus dynamics, the average subspace angle will be largely constant from prestimulus to the poststimulus period until choice and movement initiation signals begin to emerge for the fastest RTs (~ 300 ms). The subspace angle (Fig. 4D) between the first point in the prestimulus period and subsequent timepoints was < 90^*°*^ before and after checkerboard onset and only increased when movement initiation began to happen for the fastest RTs (p *<* 0.02, bootstrap). Similarly, the angle between adjacent trajectories (Fig. 4E) was largely similar throughout the trial for each direction and only begun to change after choice and movement initiation signals began to emerge, suggesting that the ordering of trajectories by RT was largely preserved well into the poststimulus period. These results imply that the initial condition was strongly predictive of the poststimulus state and eventual RT (p *<* 0.02, bootstrap), again consistent with the predictions of the dynamical systems approach.

Third, we examined if the velocity of the peristimulus dynamics was faster for faster RTs compared to slower RTs. For this purpose, we used KiNeT to find the timepoint at which the position of a trajectory is closest (minimum Euclidean distance) to the reference trajectory, which we call *Time to reference* (*t*_*ref*_, Fig. 4F). Trajectories slower than the reference trajectory will reach the minimum Euclidean distance relative to the reference trajectory later in time (i.e., longer *t*_*ref*_), whereas trajectories faster than the reference trajectory will reach these positions earlier (i.e., shorter *t*_*ref*_). Given that trajectories are compared relative to a reference trajectory, *t*_*ref*_ can thus be considered an indirect estimate of the velocity of the trajectory at each timepoint. Note, *t*_*ref*_ was referred to as speed in Remington et al. (2018b). Although a trajectory could reach the closest point to the reference trajectory later due to a slower speed, it could also be due to unrelated factors such as starting in a position in state space further from movement onset or by taking a more meandering path through state space. All of these effects are consistent with a longer *t*_*ref*_ and a slower velocity, but not necessarily a slower speed.

KiNeT revealed that faster RTs involved faster pre- and poststimulus dynamics whereas slower RTs involved slower dynamics as compared to the reference trajectory (trajectory associated with the middle RT bin, cyan) (Fig. 4F). There was also a positive correlation between RT bin and *t*_*ref*_ as measured by KiNeT at −100 ms before checkerboard onset (r = 0.82, p = 1.96 × 10^−3^). Additionally, we found that the overall scalar speed of trajectories in the prestimulus state for the first six dimensions (measured as a change in Euclidean distance over time and averaged over the 400 ms prestimulus period) covaried lawfully with RT (Fig. 4G). Thus, the ‘velocity’ of the initial condition, relative to the reference trajectory, is faster for faster RTs compared to slower RTs, coherent with the prediction of the initial condition hypothesis (Afshar et al., 2011).

Collectively, these results firmly establish that the initial condition in PMd correlates with RT and that the geometry and dynamics of these decision-related trajectories strongly depend on the position and ‘velocity’ of the initial condition consistent with the hypothesis shown in Fig. 1F (Afshar et al., 2011).

### 2.5. Initial conditions do not predict eventual choice

The previous analyses demonstrated that initial conditions strongly covaried with RT consistent with the hypothesis shown in Fig. 1F. Does the initial condition also predict choice? To investigate this issue, we first examined the covariation between prestimulus and poststimulus state with choice by measuring a choice selectivity signal identified as the Euclidean distance between the left and right choices in the first six dimensions at each timepoint. The choice selectivity signal was largely flat during the prestimulus period and increased only after stimulus onset (Fig. 4H). We also found that slower RT trials had delayed and slower increases in the choice selectivity signal compared to the faster RTs, a result consistent with the slower overall dynamics for slower compared to faster RTs (Fig. 4H). Consistent with this observation, we found a negative correlation between the average choice selectivity signal in the 125 to 375 ms period after checkerboard onset and RT (r =-0.87, p = 4.22 *×* 10^−4^).

To discriminate between the hypotheses shown in Fig. 1E, F, we interrogated the initial condition and subsequent poststimulus dynamics using a combination of single-trial analysis, decoding, and regression. We first used the Latent Factor Analysis of Dynamical Systems (LFADS) approach to estimate single-trial dynamics in a orthogonalized latent space for one of the reach choices and the easiest coherence for a single session (Pandarinath et al., 2018, 23 units). This analysis revealed that: 1) initial state for a majority of the slow RT trials are separated from the fast RT trials, 2) Initial conditions associated with a minority of the slow trials are mixed in with fast initial conditions, and 3) slower RT trajectories also appear to have more curved trajectories (Fig. 5A). All of these are consistent with the results of the trial-averaged PCA reported in Fig. 4. Finally, initial neural states related to left and right reach directions are mixed prior to stimulus onset (Fig. 5B) — again consistent with the results of the trial-averaged PCA. These single-trial dynamics suggest that prestimulus spiking activity covaries with RTs but not choice, even on single trials.

**Figure 5:**
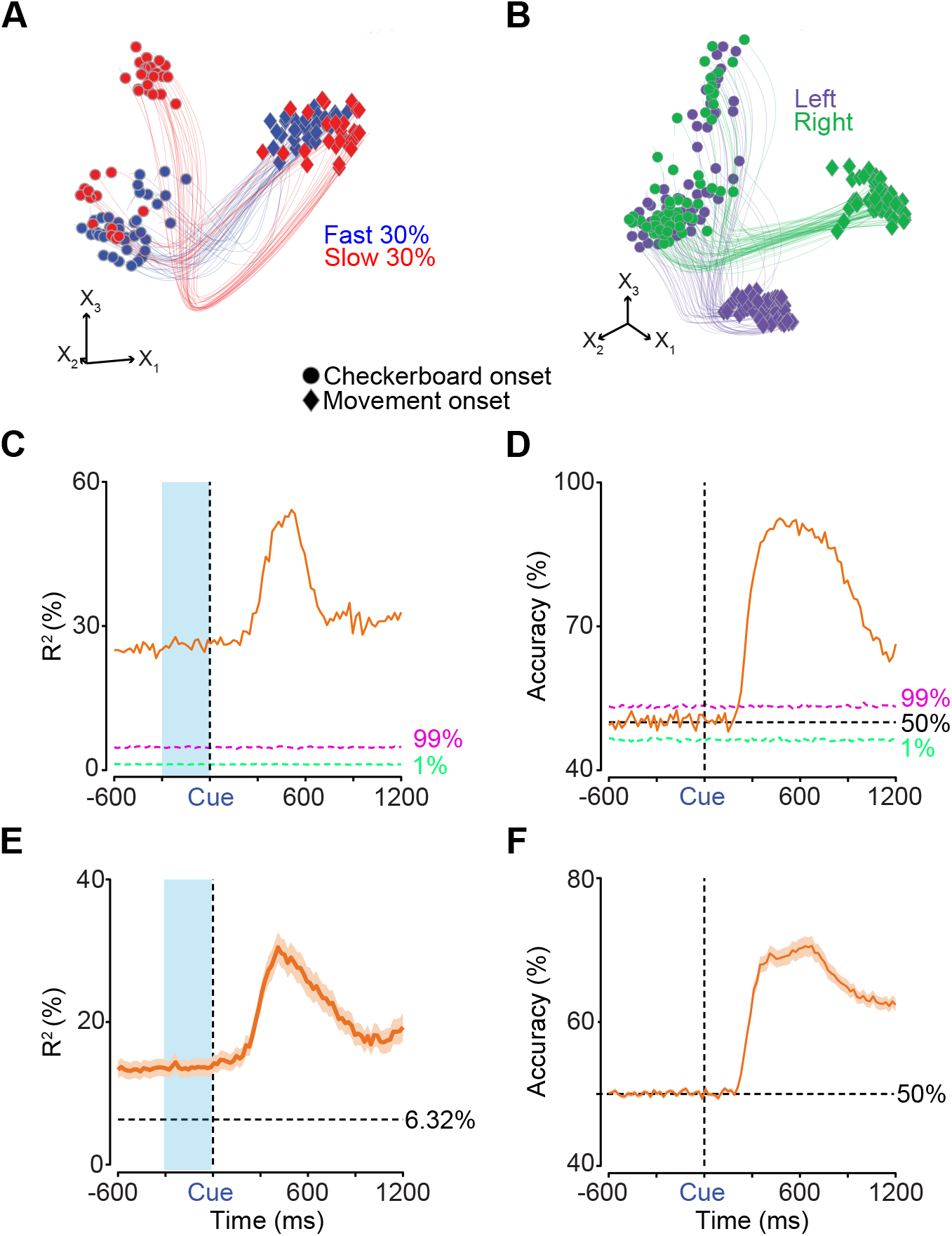
Single-trial analysis, linear regression, and decoders reveal that initial conditions predict RT but not choice. (**A/B**) LFADS (Pandarinath et al., 2018) trajectories in the space of the first three orthogonalized factors (*X*_1,2,3_), obtained via PCA on LFADS latents, plotted for (**A**) the fastest 30% of trials (blue) and the slowest 30% of trials (red) for one reach and (**B**) for left (purple) and right (green) reaches, all for the easiest coherence from a single session (23 units). Each trajectory is plotted from 200 ms before checkerboard onset (dots) to movement onset (diamonds). (**C/D**) Variance explained (*R*^2^)/decoding accuracy from linear/logistic regressions of binned spiking activity and coherence (20 ms) to predict trial-matched RTs/eventual choice from all 23 units in the LFADS session shown in **A/B**. The magenta and light green dotted lines are the 99th and 1st percentiles of *R*^2^/accuracy values calculated from averaged models of trial-shuffled (shuffled 500 times) spiking activity and RTs/choice. (**E/F**) *R*^2^/accuracy values, calculated as in **C/D**, averaged across 51 sessions. 6.32% is the average percentage of variance explained across the 51 sessions for both monkeys by regressions using stimulus coherence to predict RTs. Orange shaded area is *SEM*. 50% accuracy in **D/F** is denoted by the black dotted line.

Regression and decoding analyses of raw firing rates supported insights from the LFADS visualization (Fig. 5A) that prestimulus spiking activity would be predictive of RT. A linear regression with prestimulus spiking activity and coherence as predictors explained ~ 25% of the variance in RT from the same session used for LFADS (Fig. 5C), significantly higher than the 99th percentile of variance explained by a similar regression using trial-shuffled spiking activity instead. Identical linear regressions were performed for each of 51 sessions and *R*^2^ values were averaged across sessions. Across these sessions (Fig. 5E), prestimulus spiking activity and coherence again explained significantly more RT variance than a shuffle control of spiking activity for 47 out of 51 sessions (Mean *±* SD: 13.50 ± 8.57%, 4.70 ± 3.61%, one-tailed binomial test, p = 1.11 *×* 10^−10^, Fig. S2A).

Note, prediction of RT by spiking activity was not just an artifact of RT covarying with the coherence. A linear regression with binned spiking activity and coherence as predictors explained significantly more variance in RTs in all prestimulus bins than a linear regression of RTs with solely coherence as the predictor (only the last prestimulus bin is reported here: Mean *±* SD: 13.66 ± 8.9%, 6.32 ± 5.97%; Wilcoxon rank sum comparing median *R*^2^, p = 2.97 *×* 10^−9^, Fig. 5E). Therefore, nearly equal amounts of RT variance are explained by prestimulus neural spiking activity (*~* 7%) and the coherence of the eventual stimulus (6.32%, Fig. 5E).

In contrast, a logistic regression using binned spiking activity to predict choice, failed to predict choice, during the prestimulus period, more than the 99th percentile of accuracy from a logistic regression using trial-shuffled spiking activity (Fig. 5D). Similar logistic regressions were built for each session and accuracy was averaged across bins and sessions. The average prestimulus accuracy in predicting choice (Fig. 5F) was no better than the 99th percentile of averaged prestimulus accuracy from similar logistic regressions built on trial-shuffled spiking activity (Mean ± SD: 50.08 ± 0.51%, 50.00 ± 0.03%, only one session was larger than the shuffled data out of 51 comparisons, one-tailed binomial test, p > 0.999, Fig. S2B). Prestimulus spiking activity was no better than chance at predicting eventual choice even when trials were grouped by RT bins (Fig. S2C). These results are a key line of evidence in support of the hypothesis outlined in Fig. 1F that initial conditions covary with RT but not choice.

### 2.6. Inputs and initial conditions both contribute to the speed of poststimulus decision-related dynamics

Thus far we have shown that the initial conditions predict RT but not choice. Our monkeys clearly demonstrate choice behavior that depends on the sensory evidence, and also are generally slower for harder compared to easier checkerboards. These behavioral results and the dynamical systems approach make two key predictions: 1) sensory evidence (i.e. the input), should modulate the rate at which choice-selectivity emerges after stimulus onset and 2) the overall dynamics of the choice selectivity signal should depend on both sensory evidence and initial conditions.

To test the first prediction, we performed two analyses. First, we performed a PCA on firing rates of PMd neurons organized by stimulus coherence and choice. Fig. 6A shows the state space trajectories for the first three components. In this space, activity separates faster for easier compared to harder coherences. Consistent with this visualization, choice selectivity increases faster for easier compared to harder coherences (Fig. 6B). These results suggest that poststimulus dynamics are at least in part controlled by the sensory input consistent with the predictions of the dynamical systems hypothesis.

**Figure 6:**
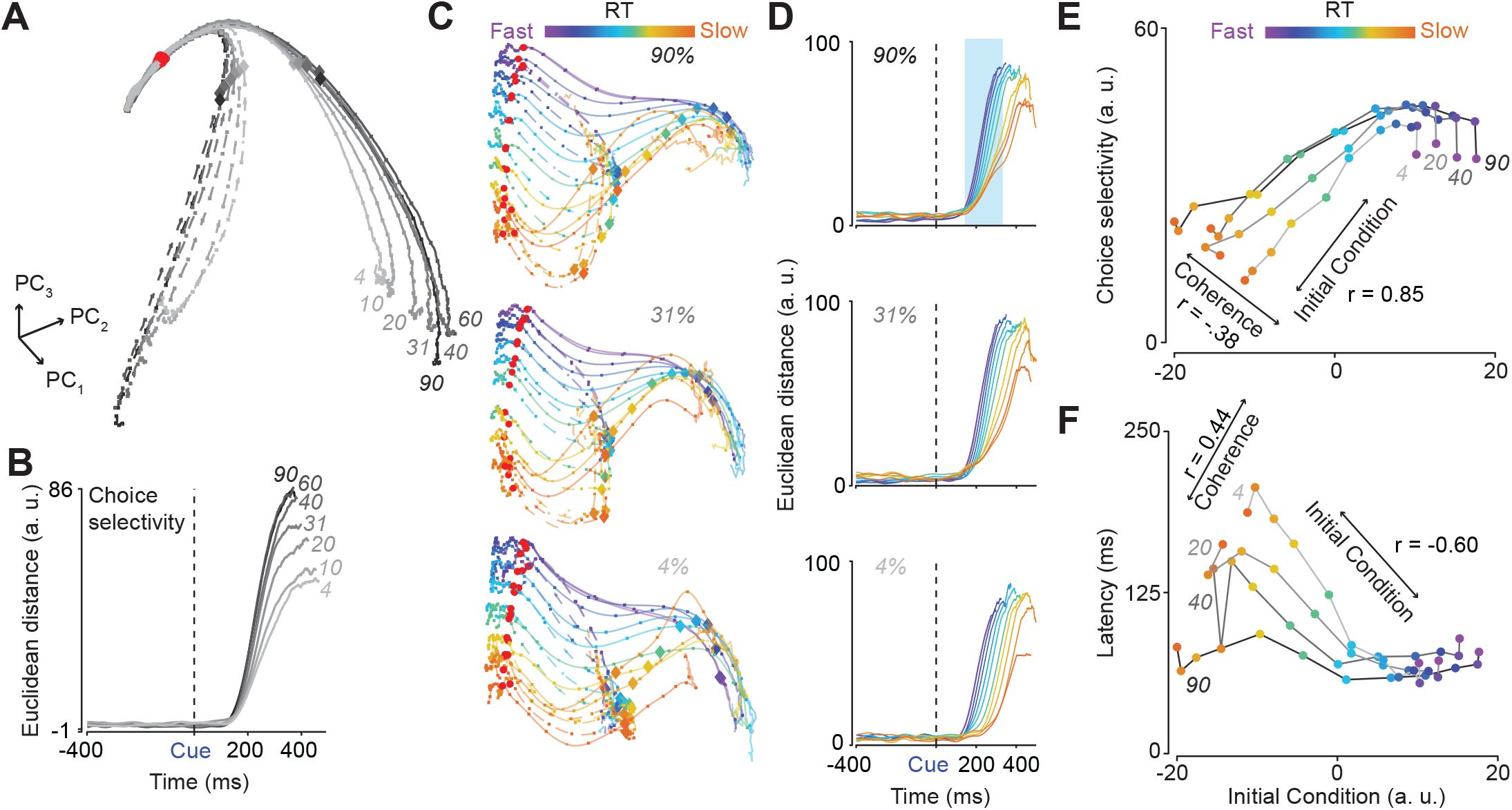
Initial conditions and inputs determine speed of dynamics and ultimately choice and RT behavior. (**A**) State space of the first three PCs (*PC*_1,2,3_) of a PCA of firing rates of all 996 units aligned to checkerboard onset (red dots) conditioned on stimulus coherence and choice. Clear poststimulus separation as a function of choice and coherence, but no observable prestimulus (−400 ms to 0 ms) separation. Diamonds and squares, color matched to their respective trajectories, indicate 250 ms post-checkerboard onset and 20 ms time steps respectively. (**B**) Choice selectivity signal measured as the Euclidean distance in the first six dimensions between left and right reaches as a function of stimulus coherence. (**C**) State space trajectories of the 1st, 2nd and 4th PCs of PCAs conditioned on RT bins and action choice within three stimulus coherences (90%, 31%, & 4%). (**D**) Choice selectivity signal for each of the three coherences shown in **C** as a function of RT bin. (**E**) Plot of the magnitude of the choice selectivity signal (averaged over the time period from 125 to 375 ms after checkerboard onset) as a function of the initial condition within each coherence. As expected easier coherences lead to higher choice selectivity signals regardless of RT, but the rates and the latencies of this signal depend on the initial condition as well as sensory evidence. (**F**) Latency of the choice selectivity signal as a function of the initial condition and for each stimulus coherence. As expected from **D**, the latency is largely flat for the easier coherences and faster RT bins (regardless of coherence), but slower for the harder coherences. For clarity, only four of the seven coherences are shown in **E & F**.

To test the second prediction of how sensory evidence and initial conditions jointly impact the speed of post-stimulus dynamics, we performed a PCA of PMd firing rates conditioned on RT and choice *within a coherence*. To obtain these trajectories, we first calculated trial-averaged firing rates for the various RT bins within each coherence. We then projected these firing rates into the first six dimensions of the PC space organized by choice and RTs (Fig. 4A & B). This projection preserved more than 90% of the variance captured by the first six dimensions of the data organized by RT bins and choice within a coherence which ranged from 75 to 80% of the total variance of the data for a given coherence. Consistent with the results in Fig. 4B, the prestimulus state again correlates with RT even within a stimulus difficulty (Fig. 6C).

To assess how inputs and initial conditions jointly influenced decision-related dynamics, we again computed the time-varying choice selectivity signal (*CS*(*t*)) by computing the high-dimensional distance between left and right trajectories at each timepoint for each of the RT bins and coherences. Fig. 6D shows this choice selectivity signal as a function of RT for the three different coherences shown in Fig. 6C. For the easiest coherence, the choice selectivity signal starts ~ 100 ms after checkerboard onset and it increases faster (i.e., steeper slope) for faster RTs compared to slower RTs (Fig. 6D, top panel, blue highlight box). In contrast, for the hardest coherence, the choice selectivity signal is more delayed for the slower RTs compared to the faster RTs, while a similar slope effect is still observed (i.e., steeper slope for fast RTs as compared to slow RTs) (Fig. 6D, bottom panel). These plots suggest that inputs and initial conditions combine and alter the rate and latency of choice-related dynamics.

We quantified these patterns by first measuring the rate at which choice-selectivity emerges. Our metric was the average choice selectivity signal in the 200 ms period from 125 to 325 ms after checkerboard onset as a function of the initial condition and for each of the 7 coherences. We obtained an estimate of the initial condition by using a PCA to project the average six-dimensional location in state space in the −300 ms to −100 ms period before checkerboard onset for each of these conditions on to a one-dimensional axis (see 4.14). As Fig. 6E shows, the rate at which the choice selectivity signal emerges is greater for easier coherences across the board but also weaker or stronger depending on the initial condition. Furthermore, when coherence is fixed, the average rate of the choice selectivity signals depends on the initial condition. A partial correlation analysis found that the rate at which choice selectivity emerges depends on both the initial condition (r = 0.85, p < 0.001) and the sensory evidence (r = −0.38, p < 0.001). These results are key evidence that choice-selective, decision-related dynamics are controlled both by the initial condition and the sensory evidence.

We also measured the latency at which choice selectivity emerged and how it depended on initial condition and sensory inputs. To estimate latency, we fit the choice selectivity signal (CS(t)) using a piecewise function as detailed in 4.13. Fig. 6F plots the latency of the choice selectivity signal (t_Latency_) as a function of the sensory input and the initial condition. Latencies are slower when the initial condition is in the slow RT state and the sensory input is weak, but faster for strong inputs and when the initial condition is in a fast RT state. Again, a partial correlation analysis found that the latency of choice selectivity depends on both the initial condition (r = −0.60, p < 0.001) and coherence (r = 0.44, p < 0.001).

Collectively, these results strongly support a dynamical system for decision-making where both initial conditions and inputs together shape the speed of decision-related dynamics and behavior, whereas poststimulus dynamics alone control choice.

### 2.7. The outcome of the previous trial influences the initial condition

So far we have demonstrated that the initial condition, as estimated by prestimulus population spiking activity, explains RT variability and poststimulus dynamics in a decision-making task. However, why initial conditions fluctuate remains unclear. One potential source of prestimulus neural variation could be post-outcome adjustment, where RTs for trials following an error are typically slower or occasionally faster than RTs in trials following a correct response (Danielmeier and Ullsperger, 2011; Purcell and Kiani, 2016; Dutilh et al., 2012).

We examined if post-outcome adjustment was present in the behavior of our monkeys. We identified all error, correct (EC) sequences and compared them to an equivalent number of correct, correct (CC) sequences. The majority of the data are from sequences of the form “CCEC” (78%), while the remainder of “EC” sequences were compared to other “CC” sequences (22%). Associated RTs were aggregated across both monkeys and sessions. We found that correct trials following an error were significantly slower than correct trials following a correct trial (M ± SD: 447 ± 117 ms, 428 ± 103 ms; Wilcoxon rank sum comparing median RTs, p = 8.44 × 10^−136^, SFig. 3A). Additionally, we found that correct trials following a correct trial were modestly faster than the correct trial that preceded it (M ± SD: 428 ± 103 ms, 431 ± 101 ms; Wilcoxon rank sum comparing median RTs, p = 8.48 × 10^−4^, SFig. 3A). Thus, trials where the previous outcome was a correct response led to a trial with a faster RT, whereas trials where the previous outcome was an error led to a trial with a slower RT.

Such changes in RT after a previous trial were mirrored by corresponding shifts in initial conditions. A PCA of trial-averaged firing rates organized by previous trial outcome and choice revealed that prestimulus population firing rate covaried with the previous trial’s outcome. Post-error correct trials, hereafter post-error trials, showed the largest prestimulus difference in firing rates as compared to other trial outcomes (Fig. 7A, B).

**Figure 7:**
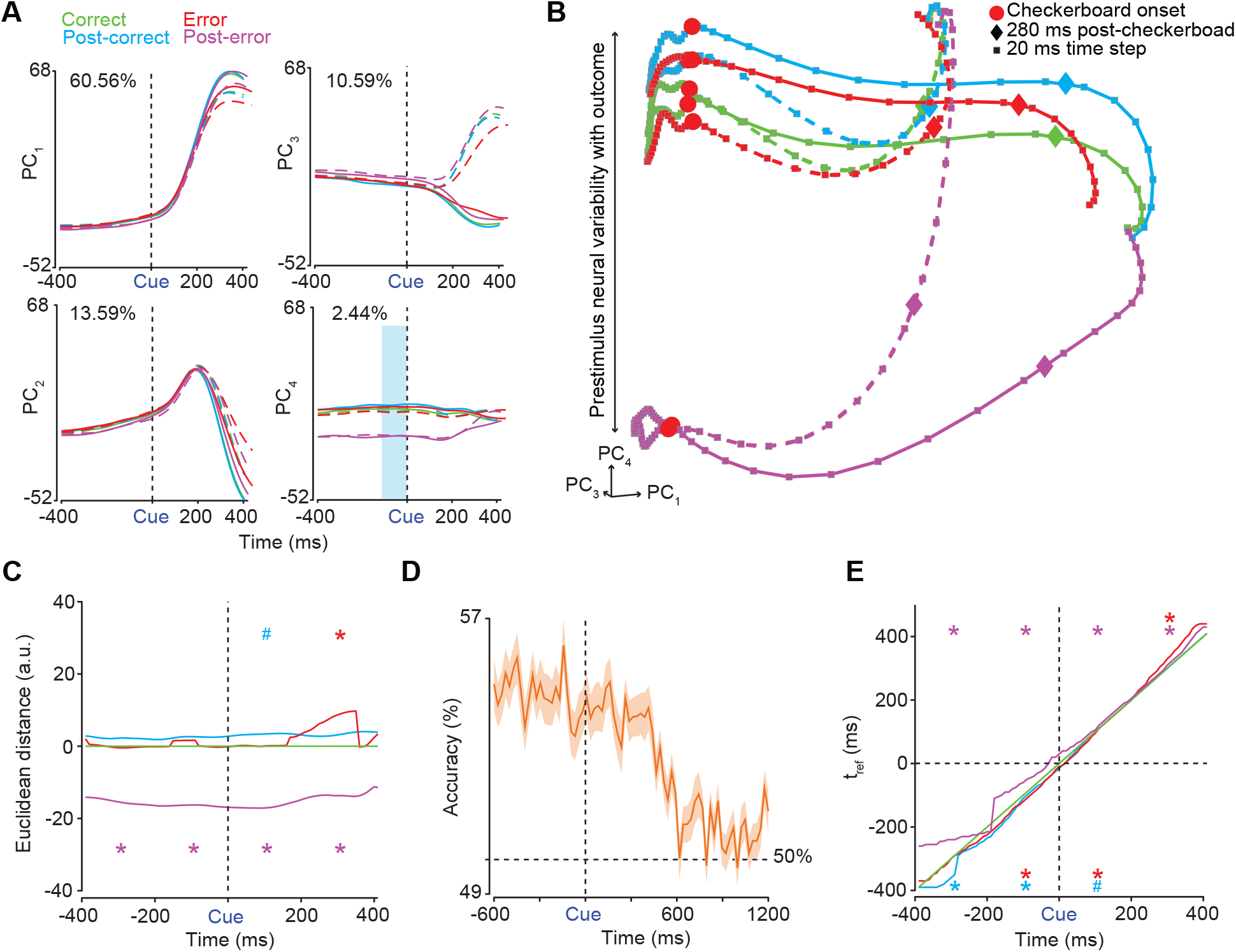
Prestimulus neural activity covaries with the previous trial’s outcome. (**A**) The first four PCs (*PC*_1,2,3,4_) of trial averaged firing rates aligned to checkerboard onset (‘Cue’ & black dashed line) of all 996 neurons from monkeys T & O and all sessions organized by choice (right - dashed lines, left - solid lines) and trial outcome (green - correct trial, cyan - correct trial following a correct trial, red - error trial, and magenta - correct trial following an error trial). Percentage variance explained by each PC presented at the top of each plot. (**B**) 1st, 3rd and 4th PC (*PC*_1,3,4_) state space aligned to checkerboard onset (red dots). Plotting of PCs extends 400 ms before checkerboard onset and 400 ms after. Observe how neural activity separates as a function of outcome, but not by choice, up to 400 ms before stimulus onset. Different colored squares and diamonds indicate 20 ms time steps and 280 ms post-checkerboard onset respectively. (**C**) “KiNeT distance” analysis demonstrating that trajectories are spatially organized with post-error trials furthest from other trial types peri-stimulus as compared to a reference trajectory (green, middle trajectory). (**D**) Accuracy of logistic regression of spiking activity from the current trial used to predict the outcome of the previous trial. Orange outline is SEM. (**E**) “KiNeT Time to reference” (*t*_*ref*_) analysis reveals that prestimulus ‘velocity’ is slower for post-error trials as compared to the reference trajectory (green, middle trajectory). In **C & E** the x-axis is labelled “Time (ms)”, this should be understood as time on the reference trajectory. *Abbreviation*: a.u. - arbitrary units, * - p < 0.05, # - p = 0.05.

A KiNeT analysis (Remington et al., 2018b) further corroborated prestimulus firing rate covariation with the previous trial’s outcome. Peri-stimulus trajectories for post-error trials occupied the reflected side of state space, relative to the reference trajectory (“Correct” trials), as compared to all other trial types (Fig. 7C). The averaged, windowed (i.e., −400:−200 ms, −200:0 ms, 0:200 ms, 200:400 ms) post-error trajectory was significantly different, at p< 0.05, from equivalently averaged shuffled data (Fig. 7C). In a similar finding a decoder revealed that the current trial’s spiking activity can predict, at greater than chance levels, the previous trial’s outcome from before stimulus onset until about the overall mean RT, ~ 450 ms (equal numbers of correct and error trials, were used in training the decoder, Fig. 7D), suggesting that the previous trial’s outcome has an effect on the current trial’s pre- and poststimulus population firing rates.

KiNeT analyses suggested that post-error trials also had significantly slower prestimulus trajectories as compared to the reference trajectory, p < 0.05 for all prestimulus windows (i.e., −400:-200 ms & −200:0 ms), (Fig. 7E), suggesting that error trials or, similarly, infrequent outcomes (Danielmeier and Ullsperger, 2011) result in slower population dynamics in the following trial. Additionally, trials that follow correct trials (errors generally followed correct trials) have slightly faster prestimulus dynamics as compared to the reference trajectory, p < 0.05 for both prestimulus windows for post-correct trials and just one prestimulus window for error trials (−200:0 ms) (Fig. 7E). Finally, error and post-error trials have the slowest poststimulus trajectories in the last poststimulus window (200:400 ms), (p<0.05, Fig. 7E) consistent with their longer RTs. Altogether, these results complement behavioral results in that the initial condition shifts as a function of previous trial outcome and not just due to errors. These results suggest that slower or faster RTs after an error or correct trial are at least partially due to slower or faster prestimulus dynamics respectively (see SFig. 3B for complementary findings in single trials).

These results strongly suggest that some of the initial condition covariation with RT Fig. 4 might be related to the previous trial’s outcome. To test this hypothesis, we performed two analyses: First, we wanted to know how much of the variance of the firing rate data organized by RT and choice could be accounted for by the subspace spanned by the first six dimensions of the PCA organized by previous trial’s outcome and choice (Elsayed et al., 2016, “outcome subspace”, Fig. 7A, B). We chose the first six dimensions (explains ~ 90% of the variance) of this outcome subspace as these dimensions were significantly above or equal to noise components (SFig. 3C). This analysis revealed that 77.19% of the total variance for the firing rates organized by RT and choice was explained by the first six dimensions of the outcome subspace suggesting that the previous trial’s outcome has a large impact in explaining prestimulus firing rate covariation with RT.

In a parallel analysis we performed a dPCA (Kobak et al., 2016) on the population firing rates in the 600 ms before checkerboard onset organized by previous trial’s outcome and choice, and another organized by RT and choice. The respective axes that maximally separated as a function of previous trial’s outcome and that maximally separated as a function of RT demonstrated significant overlap with an angle of 47.8^*°*^ between them. These results suggest that the previous trial’s outcome leads to a shift in prestimulus dynamics consistent with determining the speed of the dynamics and therefore eventual RTs.

Lastly, we examined if there were differences in pre- and poststimulus state with respect to choice between the different trial outcomes. Again the high-dimensional Euclidean distance between left and right choice trajectories was largely flat during the prestimulus period and increased only after stimulus onset for all trial outcomes (SFig. 3D). We also found that the separation between choices increased slower for error trials as compared to all other trial outcomes (SFig. 3D).

These findings are consistent with the dynamical systems approach as they demonstrate that initial condition before stimulus onset is dependent upon trial history and that pre- and poststimulus dynamics slow down after errors as compared to after correct trials. Collectively, the past trial’s outcome leads to different initial conditions, slower pre- and poststimulus dynamics and ultimately leads to RT variability, all in line with the hypothesis in Fig. 1F.

## 3. Discussion

Our goal in this study was to rigorously identify a dynamical system for the neural population activity underlying decision-making as recently demonstrated in studies of neural population dynamics related to motor planning and timing (Afshar et al., 2011; Remington et al., 2018b; Shenoy et al., 2013; Vyas et al., 2020a). To this end, we investigated the neural population dynamics in PMd of monkeys performing a red-green RT decision-making task (Chandrasekaran et al., 2017; Coallier et al., 2015). The prestimulus neural state in PMd, proxy for the initial condition of the dynamical system, was strongly predictive of RT, but not choice. We observed these effects across and within stimulus difficulties and also on single trials. Furthermore, faster RT trials had faster neural dynamics and separate initial conditions from slower RT trials. Additionally, poststimulus, choice-related dynamics were altered by the inputs with easier checkerboards leading to faster dynamics than harder ones. Finally, these initial conditions and the behavior for a trial depended on the previous trial’s outcome, where RTs and prestimulus trajectories were slower for post-error compared to post-correct trials. Together, these results suggest that decision-related neural population dynamics in PMd can be well described by a dynamical system where the speed of the choice (the output of the system) is strongly set by its initial conditions. However, the eventual choice itself is determined by the input and the speed of these choice-related dynamics depends on the initial condition. Finally, the outcome of the trial affects the initial condition of the next trial.

At the highest level, these observations are another compelling demonstration of the power of the dynamical systems approach (alternatively, “computation through dynamics”) to explain the link between the time-varying activity of neural populations and behavior (He, 2013; Vyas et al., 2020b; Briggman et al., 2005; Chaisangmongkon et al., 2017; Mante et al., 2013; Mazor and Laurent, 2005; Remington et al., 2018b; Stroud et al., 2018). Regardless of species or brain region, an increasingly common finding is that neurons associated with cognition and motor control are often heterogeneous and demonstrate complex time-varying patterns of firing rates and mixed selectivity (Chaisangmongkon et al., 2017; Rigotti et al., 2013; Mante et al., 2013; Machens et al., 2010; Hanks et al., 2015). Simple models or indices although attractive to define are often insufficient to summarize the activity of these neural populations (Chandrasekaran et al., 2018; Chaisangmongkon et al., 2017; Mante et al., 2013), and even if one performs explicit model selection on single neurons using specialized models (Latimer et al., 2015), the results can be brittle because of the heterogeneity inherent in these brain regions (Chandrasekaran et al., 2018). The dynamical systems approach addresses this problem by using dimensionality reduction and optimization techniques to understand collective neuronal activity of different brain regions and tasks, generally summarizing large population datasets in orders of magnitude fewer dimensions than were recorded from (Okazawa et al., 2021; Mante et al., 2013; Machens et al., 2010). Here, we demonstrated that >90% of the variance from the firing rate activity of nearly 1,000 neurons in PMd during decisions could be explained in just a few (six) dimensions.

Besides providing a compact description of population activity, there are three other clear advances afforded by using a dynamical systems approach to study decisions. First, we find lawful relationships between the low-dimensional activity of neural populations and task variables such as choice, RT, stimulus difficulty and past outcomes (Mante et al., 2013; Okazawa et al., 2021). Second, this lawful relationship can be understood as emerging from a dynamical system that is parameterized by initial conditions and inputs that subsumes much of decision-making behavior (Vyas et al., 2020a). Finally, this dynamical system naturally bridges previously disparate findings from SAT (Murphy et al., 2016; Heitz and Schall, 2012; Bogacz et al., 2010), post-outcome adjustment (Purcell and Kiani, 2016; van den Brink et al., 2014), motor planning (Afshar et al., 2011) and timing (Remington et al., 2018b) and provides a common framework for deriving models for the neural computations underlying decision-making.

In the remainder of the discussion, we further discuss the implications of our identified dynamical system for decision-making models, and unpack the factors that may underlie initial conditions.

Our results are an important and significant advance over a previous study of dynamics in PMd during reach planning (Afshar et al., 2011). As described previously in this study, Afshar et al. (2011) showed that in a delayed reach task, the position and velocity of the initial conditions correlated with RT. However, it was unclear from the study, what the role of inputs was and how changes in initial conditions emerge across trials. Our study answers both questions and provides a clear account of how both initial conditions and inputs jointly control the dynamics in PMd, a key brain region involved in mapping sensory cues to actions (Kurata and Hoffman, 1994). The sensory evidence, which acts as the input combines with initial conditions determining the choice of the monkeys and also alters the speed of the choice. We also demonstrated that changes in initial conditions emerge due to the outcome of the previous trial with errors leading to large shifts in the initial condition and significantly altering subsequent dynamics.

Our results, mainly that decision-related neural activity and behavior are well described by a dynamical system dependent upon both initial conditions and inputs, are inconsistent with simple drift diffusion models (DDMs) where decision-making behavior is solely driven to a bound by accumulation of sensory evidence (Ratcliff, 1978; Ratcliff et al., 2016; Hawkins et al., 2015). Including variable drift rates and starting points in a DDM would be insufficient towards recapitulating prestimulus decision-related signals that covary with RT. Variable non-decision times could potentially explain the RT behavior reported here. However, the neural effect of a change in non-decision time is thought to relate to changes in the initial latency of decision-related responses and does not predict changes in the prestimulus neural state. Thus, while simple DDMs with a variable non decision time may explain the behavior observed herein they would fail to recreate the observed variability in the initial condition.

We believe that cognitive process models with an additive or multiplicative stimulus-independent gain signal, previously described as “urgency” and successfully used to describe monkey behavior and neural activity (Cisek et al., 2009; Thura et al., 2014; Cowley et al., 2020; Murphy et al., 2016), could faithfully model the behavior *and* the neural dynamics. A variable additive gain signal, which adds inputs to accumulators for left and right choices in a race model for decisions, would lead to different initial conditions and thus faster dynamics for faster RTs and slower dynamics for slower RTs (Murphy et al., 2016). Similarly, a multiplicative gain signal would also lead to differences in both the initial firing rates and control the speed of decision-making behavior (Murphy et al., 2016; Cisek et al., 2009). Both types of gain signals generate similar predictions about RT and choice behavior and are often difficult to distinguish using trial-averaged firing rates as done here. One way to resolve this impasse would be to employ single-trial analysis (Peixoto et al., 2021) of neural responses in multiple brain areas using a task paradigm that dispenses sensory evidence over the course of a trial such as in the tokens (Thura and Cisek, 2014) or pulses task (Hanks et al., 2015).

Typically, researchers have focused on the slowing down of responses after an error, a phenomenon termed post-error slowing (Dutilh et al., 2012; Purcell and Kiani, 2016). However, our findings suggest that both correct and error outcomes can influence the pre- and poststimulus decision-making neural dynamics on subsequent trials suggesting that post-error slowing could be better understood under the umbrella of post-outcome adjustments (Danielmeier and Ullsperger, 2011). It is currently unclear how these post-outcome adjustments in PMd emerge. One possibility is that these adjustments emerge from the internal dynamics of PMd itself. Errors vs. correct trials could lead to a shift in the initial condition due to recurrent dynamics that occur in PMd due to the presence or absence of reward. Such error related signals have been observed in premotor and motor cortex and have even been used to augment brain computer interfaces (Even-Chen et al., 2017). Alternatively, the changes observed in PMd could emerge from inputs from other brain areas such as the anterior cingulate cortex (ACC) which is known to monitor trial outcome (Hyman et al., 2013), or the supplementary motor area (SMA), which has been implicated in timing of motor actions and evaluative signals related to outcome (Bogacz et al., 2010; Ullsperger et al., 2014). Simultaneous recordings in PMd and these brain areas are necessary to tease apart the contribution, if any, of these areas to the initial condition changes observed in PMd.

Barring Fig. 5 and Fig. S3, our description of decision-related dynamics largely focused on trial-averaged activity. Even with such a constraint we were able to identify that the position of initial conditions predict RT, and are modified by the outcome of the previous trial and that the dynamics for faster RT trials are further along the movement initiation path compared to slower RT trials. We also demonstrated that sensory inputs combined with initial conditions to alter the speed of dynamics and drive choice-related behavior. We believe that even further insights will be available using single-trial analysis. In particular, here we were unable to fully characterize the relative contributions of the position of the initial condition and the velocity of the initial condition to decision-related dynamics and behavior. We anticipate that further analyses of the curvature, velocity relative to the mean trajectory, path length, and speed of the trajectories will lead to an even better description of the single-trial dynamics underlying decisions as has been done for motor planning (Afshar et al., 2011). Note, we were unable to fully perform such analyses in the current study as we often had only a few neurons per session in Monkey O. The session shown in Fig. 5 and Fig. S3 was an exception as we had 23 well-modulated units in Monkey T.

We have shown that the outcome of the previous trial alters the initial conditions for subsequent trials. There are certainly other factors that lead to changes in the initial conditions. In particular, recent studies have shown that both neural activity and behavior as indexed by RT, performance, and pupil size drifts over slow time scales and that these slowly drifting signals are likely a process independent of deliberation on sensory evidence (Cowley et al., 2020; Ferguson and Cardin, 2020). Such effects often emerge over several hours. We believe that such effects could also contribute to the changing initial conditions observed in our study. However, we were unable to assess these effects as 1) we did not measure pupil size, 2) significant amounts of our data were collected with single electrode recordings over short time periods (often 10-15 minutes or so for a tranche of 300-500 trials), and 3) even in sessions where Plexon U-probes were used to simultaneously record from neural populations we often paused the task whenever the animal disengaged from the task or had a sudden decrease in performance. Furthermore, after such pauses we generally increased reward sizes to remotivate the animals. These interventions are often standard for electrophysiological recordings in behaving monkeys but preclude the assessment of the effects of slow fluctuations on decision-making. Nevertheless, we believe that such effects are likely to be an additional crucial source of variability for the initial condition, especially given that it was found to be a factor independent of sensory evidence (Cowley et al., 2020) as in our study, and likely alters decision-making dynamics and behavior. A rich area for future research is to assess whether the same effects observed in V4, and caudal prefrontal cortex in Cowley et al. (2020) also occurs for perceptual decisions in PMd.

We found that prestimulus neural activity in PMd and in this task did not covary with or predict eventual choice. However prestimulus neural activity in lateral intraparietal cortex was found to be predictive of choice for low coherence or harder random dot stimuli (Shadlen and Newsome, 2001). Our lack of an observed covariation between the initial condition and choice may be due to the randomization of target configurations, thus the monkeys in our experiment were disincentivized from preplanning a reach direction. To be clear, our lack of a finding does not preclude prestimulus activity in other brain areas or even in PMd with different tasks from covarying with choice (Peixoto et al., 2018).

We believe that the effects we see where the initial conditions predict the RT of the animal in a cognitive task are likely to be observed in many brain areas. For example, previous results recorded in monkey dorsomedial prefrontal cortex during timing tasks (Remington et al., 2018b) and in motor cortex/PMd from motor planning tasks (Afshar et al., 2011) bear out the contention that our observation of prestimulus PMd neural population activity covarying with and predicting RTs in a decision-making task is likely not solely localized to PMd or constrained to occur only in this task. In fact, differences in baseline modulation of neural activity between speed and accuracy conditions of speed-accuracy tradeoff tasks (Heitz and Schall, 2012) is found in frontal eye field (Heitz and Schall, 2012) and pre-supplementary motor area (Bogacz et al., 2010). We also showed that prestimulus beta band activity in this same task was correlated with RT (Chandrasekaran et al., 2019). Additionally, in a study of post-error slowing the level of prestimulus phase synchrony in fronto-central electrodes, was found to positively correlate with the speed of RTs (van den Brink et al., 2014). These findings of neural activity changing as a result of different conditions of a speed-accuracy tradeoff task or being predictive of RTs, strongly suggest that initial conditions in multiple brain regions, and potentially some putative fronto-central motor network, effect the speed of a response. In other words, changes in the initial conditions in various brain regions before stimulus onset is likely not a localized effect and suggests either broad signalling (Derosiere et al., 2022) from some source or even feed-forward/feedback mechanisms between brain regions.

### 3.1. Conclusion

Research employing dynamical systems approaches demonstrate that future population level activity and behavior is sensitive to initial conditions such that initial conditions were predictive of RTs in motor planning or timing tasks (Afshar et al., 2011; Remington et al., 2018b). However it was unclear whether decision-related neural activity was similarly sensitive to initial conditions and if so, how such sensitivity might interact with sensory evidence accumulation, a well-studied aspect of decision-making (e.g., Roitman and Shadlen, 2002). Our first main contribution is that we observe prestimulus neural dynamics predictive of the RT of a decision, *equivalent to the predictive power of the eventual stimulus itself*, despite lacking an explicit manipulation of speed-accuracy tradeoff. Our second main contribution was to show that both initial conditions and sensory evidence influenced choice-related neural population dynamics and ultimately behavior. Finally, our third contribution was to show that initial conditions depended on previous outcomes, and, in turn, altered poststimulus dynamics and RTs. We believe that this suite of findings through the lens of the dynamical systems approach is a starting point for understanding the dynamical system underlying decision-making behavior. The insights from this study could be further expanded via single-trial analysis of simultaneous recordings in multiple decision-related regions, by examining how baseline neural activity predicts various aspects of behavior, and ultimately how behavior or global state then feeds back into initial conditions.

## 4. Methods

Several method sections are adapted from Chandrasekaran et al. (2017) as the same data set is reanalyzed in this study. For completeness and readability, some aspects are replicated here, but much of the methods focuses on key details about the various dimensionality reduction techniques such as PCA, decoding, and LFADS analyses.

### 4.1. Code and data availability

MATLAB scripts for generating all the figures are available with the paper along with the relevant data. HTML code that allows free rotation of the trajectories in principal component (PC) spaces are also available in the ZIP file.

### 4.2. Subjects

Experiments were performed using two adult male macaque monkeys (*Macaca Mulatta*; monkey T, 7 years, 14 kg & monkey O, 11 years, 15.5 kg) trained to touch visual targets for a juice reward. Monkeys were housed in a social vivarium with a normal day/night cycle. Protocols for the experiment were approved by the Stanford University Institutional Animal Care and Use Committee. Animals were initially trained to come out of their housing and to sit comfortably in a chair. After initial training (as described in Chandrasekaran et al. (2017)), monkeys underwent sterile surgery where cylindrical head restraint holders (Crist Instrument Co., Inc., Hagerstown, MD, United States) and standard circular recording cylinders (19 mm diameter, Crist Instrument Co., Inc.) were implanted. Cylinders were placed surface normal to the cortex and were centered over caudal dorsal premotor cortex (PMdc; +16, 15 stereotaxic coordinates, see Fig. 2G). The skull within the cylinder was covered with a thin layer of dental acrylic.

### 4.3. Apparatus

Monkeys sat in a customized chair (Synder Chair System, Crist Instrument Co., Inc.) with their head restrained. The arm that was not used to respond in the task was gently restrained with a tube and cloth sling. Experiments were controlled and data collected using a custom computer control system (Mathworks’ xPC target and Psychophysics Toolbox, The Mathworks, Inc., Natick, MA, United States). Stimuli were displayed on an Acer HN2741 monitor approximately 30 cm from the monkey. A photodetector (Thorlabs PD360A, Thorlabs, Inc., Newton, NJ, United States) was used to record the onset of the visual stimulus at a 1 ms resolution. A small reflective spherical bead (11.5 mm, NDI passive spheres, Northern Digital, Inc., Waterloo, ON, Canada) was taped to the middle finger, 1 cm from the tip, of the active arm of each monkey; right for T and left for O. The bead was tracked optically in the infrared range (60 Hz, 0.35 mm root mean square accuracy; Polaris system, NDI). Eye position was tracked using an overhead infrared camera with an estimated accuracy of 1° (ISCAN ETL-200 Primate Eye Tracking Laboratory, ISCAN, Inc., Woburn, MA, United States). To get a stable image for the eye tracking camera, an infrared mirror (Thorlabs, Inc.) transparent to visible light was positioned at a 45° angle (facing upward) immediately in front of the nose. This reflected the image of the eye in the infrared range while allowing visible light to pass through. A visor placed around the chair prevented the monkey from touching the juice reward tube, infrared mirror, or bringing the bead to its mouth.

### 4.4. Task

Experiments were made up of a sequence of trials that each lasted a few seconds. Successful trials resulted in a juice reward whereas failed trials led to a time-out of 2-4 s. A trial started when a monkey held its free hand on a central circular cue (radius = 12 mm) and fixated on a small white cross (diameter = 6 mm) for ~ 300-485 ms. Then two isoluminant targets, one red and one green, appeared 100 mm to the left and right of the central hold cue. Targets were randomly placed such that the red target was either on the right or the left trial-to-trial, with the green target opposite the red one. In this way color was not tied to reach direction. Following an additional center hold period (400-1000 ms) a static checkerboard stimulus (15 × 15 grid of squares; 225 in total, each square: 2.5 mm x 2.5 mm) composed of isoluminant red and green squares appeared superimposed upon the fixation cross. The monkey’s task was to move their hand from the center hold and touch the target that matched the dominant color of the checkerboard stimulus for a minimum of 200 ms (for full trial sequence see Fig. 2B). For example, if the checkerboard stimulus was composed of more red squares than green squares the monkey had to touch the red target in order to have a successful trial. Monkeys were free to respond to the stimulus as quickly or slowly, within an ample *~* 2*s* time frame, as they ‘chose’. There was no delayed feedback therefore a juice reward was provided immediately following a successful trial (Roitman and Shadlen, 2002). An error trial or miss led to a timeout until the onset of the next trial.

The checkerboard stimulus was parameterized at 14 levels of red (*R*) and complementing green (*G*) squares ranging from nearly all red (214 *R*, 11 *G*) to all green squares (11 *R*, 214 *G*) (for example stimuli see Fig. 2C). These 14 levels are referred to as signed coherence (*SC*), defined as 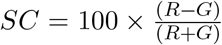 (R: 4%:90%, G:-4%:-90%). Correspondingly there are seven levels of color coherence, agnostic to the dominant color, defined as 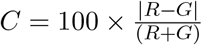 (4-90%).

The hold duration between the onset of the color targets and onset of the checkerboard stimulus was randomly chosen from a uniform distribution from 400-1000 ms for monkey T and from an exponential distribution for monkey O from 400-900 ms. Monkey O attempted to anticipate the checkerboard stimulus therefore an exponential distribution was chosen to minimize predictability.

### 4.5. Effects of coherence on accuracy and reaction time (RT)

Behavior was analyzed by fitting psychometric and RT curves on a per-session basis and averaging the results across sessions. Behavioral data was analyzed in the same sessions as the electrophysiological data. In total there were 75 sessions for monkey T (128,774 trials) and 66 sessions for monkey O (108,365 trials). On average there were ~ 1,500 trials/session. Both incorrect and correct trials for each *SC* were included for estimating RT/session.

Data were fit to a psychometric curve to characterize how discrimination accuracy changed as a function of stimulus coherence. For each session a monkey’s sensitivity to the checkerboard stimulus was estimated by estimating the probability (*p*) of a correct choice as a function of the color coherence of the checkerboard stimulus (*c*). The accuracy function was fit using a Weibull cumulative distribution function.

**Weibull cumulative distribution function:**

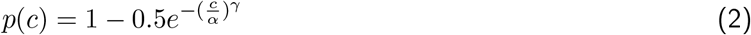

The discrimination threshold *α* is the color coherence level at which the monkey would make 81.6% correct choices. The parameter *γ* describes the slope of the psychometric function. Threshold and slope parameters were fit per session and averaged across sessions. We report the mean and standard deviation of threshold and *R*^2^ values from the fit in the text.

Mean RT was calculated per *SC* on a session-by-session basis and averaged across sessions. Results are displayed in Fig. 2E with error bars denoting 2 × *SEM* and lines between the averages to guide the eyes. RT was also regressed with *log*_10_(*C*) per session. The fit coherence-RT model was used to predict RTs and calculate *R*^2^ on a per session basis. *R*^2^ values were averaged across sessions per monkey and are reported in Fig. 2F as percentage of variance explained. The general framework and equations for linear regression and *R*^2^ calculations are provided in 4.16.

### 4.6. Electrophysiological recordings

Electrophysiological recordings were guided by stereotaxic coordinates, known response properties of PMd, and neural responses to muscle palpation. Recordings were made anterior to the central sulcus, lateral to the precentral dimple and lateral to the spur of the arcuate sulcus. Electrodes were placed in the PMd contralateral to the dominant hand of the monkey (T: right arm, O: left arm). Recording chambers were placed surface normal to the cortex to align with the skull of the monkey and recordings were performed orthogonal to the surface of the brain. Estimates of upper and lower arm representation was confirmed with repeated palpation at a large number of sites to identify muscle groups associated with the sites.

Single electrode recording techniques were used for a subset of the electrophysiological recordings. Small burr holes in the skull were made using handheld drills (DePuy Synthes 2.7 to 3.2 mm diameter). A Narishige drive (Narishige International USA, Inc., Amityville, NY, United States) with a blunt guide tube was placed in contact with the dura. Sharp FHC electrodes (*>* 6 MΩ, UEWLGCSEEN1E, FHC, Inc., Bowdoin, ME, United States) penetrated the dura and every effort was made to isolate, track, and stably record from single neurons.

180 µm thick 16-electrode linear multi-contact electrode (U-probe, see Fig. 2G; Plexon, Inc., Dallas, TX, United States); interelectrode spacing: 150 µm, contact impedance: ~ 100 kΩ) recordings were performed similarly to single electrode recordings with some modifications. Scraping away any overlying tissue on the dura, under anesthesia, and a slightly sharpened guide tube aided in slow U-probe penetration (~2-5 µm/s). U-probe penetration was stopped once a reasonable sample of neurons was acquired, potentially spanning multiple cortical layers. Neural responses were allowed to stabilize for 45-60 minutes before normal experimentation began. Monkey T had better recording yields on average (~16 units/session) than monkey O (~9 units/session). Additionally, lowering the electrode necessitated careful observation to ensure the electrode did not bend or break at the tip, or excessively dimple the dura. Therefore, it was not possible to precisely localize the U-probes with a grid system between sessions.

### 4.7. Unit selection and classification

The electrophysiological recordings consist of 996 units (546 units in T and 450 units in O, including both single neurons and multi-units) recorded from PMd of the two monkeys as they performed the task over 141 sessions. Chosen units were included as they were well isolated from other units/separated from noise and modulated activity in at least one task epoch.

U-probes were useful for recording from isolated single neurons as U-probes are low impedance (~100 kΩ) with a small contact area. A conservative threshold was used to maximize the number of well defined waveforms and to minimize contamination from spurious non-neural events. Single neurons were delineated online by the ‘hoops’ tool of the Cerebus system software client (Blackrock Microsystems, Salt Lake City, UT, United States) after the electrodes had been in place for 30 - 45 minutes. When a spike was detected via thresholding, a 1.6 ms snippet was stored and used for subsequent evaluation of the clusters as well as modifications needed for spike sorting.

Some electrodes in U-probe recordings captured mixtures of 2 or more neurons, well separated from each other and noise. In the majority of cases the waveforms were separable and labeled as single units. These separations were verified by viewing the waveforms in principal component (PC) space using custom code in MATLAB (The MathWorks, Inc., Natick, MA, United States). MatClust the MATLAB based clustering toolbox or Plexon Offline Sorter (Plexon, Inc.) were used to adjust the clusters that were isolated online.

Recording activity labeled as ‘multi-units’ were mixtures of 2 or more neurons not separable using a PCs method or consisted of recordings with waveforms only weakly separable from noise.

The number of interspike interval (ISI) violations after clustering and sorting was used to mitigate subjectivity in the classification of units. A unit was labeled as a single neuron if the percentage of ISI violations (refractory period of ≤ 1.5 ms) was ≤ 1.5%, otherwise it was labeled as a multi-unit. 801/996 PMd units were labeled as single neurons (T: 417, O: 384, median ISI violation = 0.28%, mean ISI violation = 0.43%, ~0.13 additional spikes/trial). Therefore 195/996 units were labeled as multi-unit (T: 129, O: 66, mean ISI violation = 3.36%, *~*1.4 additional spikes/trial).

Units from both monkeys were pooled together as the electrophysiological characteristics were similar. Change-of-mind trials (~2-3%) were excluded from averaging as the change in reach direction mid-movement execution made the assignment of choice ambiguous. Incorrect and correct trials arranged by choice were averaged together.

### 4.8. Peri-event firing rates

We estimated the peri-event time histograms aligned to various of events such as checkerboard onset (e.g., in Fig. 3) and for principal component analysis using the following procedure. 1) We first binned spike times for each trial at 1 ms resolution for a condition of interest (say a fast RT bin and left reaches) aligned to checkerboard or movement onset. 2) We then convolved the spike train with a Gaussian kernel (*σ* = 30 ms) to estimate the instantaneous firing rate (e.g., *r*_*i*_(*t, RT, left*)) for a trial. 3) We then used these trials to estimate the mean and standard error of the firing rate for a condition (e.g., 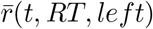).

When firing rates were aligned to checkerboard onset, we removed all spikes 50 ms before movement onset until the end of the trial. We performed this operation to ensure movement related spiking activity did not spuriously lead to ramping in the checkerboard period.

### 4.9. Principal component analysis (PCA) of PMd firing rates

PCA was used to examine firing rate variance in the recorded PMd neural population. PCA reveals dimensions that explain a large percentage of the data while making few assumptions about the underlying structure of the data. The dimensions extracted by PCA may not always be meaningful however they often align well with behavioral variables.

The general procedure for performing a PCA involved creating a 4D matrix of all 996 units and their average firing rate activity (trial spike times censored post RT trial-by-trial and convolved with 30 ms wide Gaussian kernel as explained above) windowed about checkerboard onset (~-600 ms: ~1200 ms) and organized by level of condition (e.g., coherence, RT, or past outcome) within a reach direction. Typical matrix organization was windowed firing rate x units x reach x coherence/RT/past outcome (~1800 × 996 × 2 × 7/11/2). The raw data was centered by subtracting the mean of each column (i.e. units) and then normalized by dividing by the square root of the 99th percentile of that column (i.e., soft normalization). Soft normalization reduces the bias of units with high firing rates and ensures that each unit has roughly the same overall variability across conditions. Eigenvectors, eigenvalues, and the projected data were calculated using the pca function in MATLAB.

### 4.10. Estimation of number of dimensions to explain the data

We used the approach developed by Machens et al. (2010) to estimate the number of dimensions that best described our data. The assumption of this method is that the firing rates of the *k*^*th*^ neuron for the *i*^*th*^ trial given a RT bin and choice 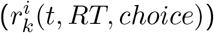 are assumed to be composed of a mean “signal” rate (*q*_*k*_(*t* | *RT, choice*)) and a “noise” rate that fluctuates across trials 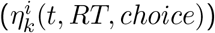.

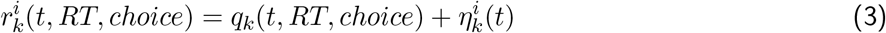

Noise here encompasses both contributions from the random nature of spike trains as well as systematic but unknown sources of variability. Averaging over trials:

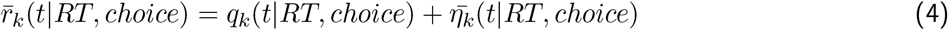

Where 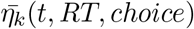 is the average noise over N instantiations (i.e., trials) of the noise term 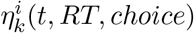.

The overall mean firing rate over time and conditions 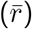 is given as:

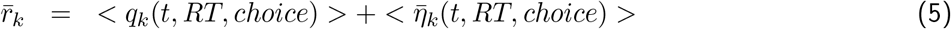

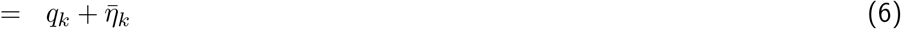

Note, none of these assumptions are strictly true. Noise may not be additive and it may depend on RT bin and may increase or decrease during various phases of the trial. However, these assumptions illustrate the problem encountered in identifying the number of dimensions to best describe the data.

Under these assumptions PCA attempts to identify a covariance matrix as

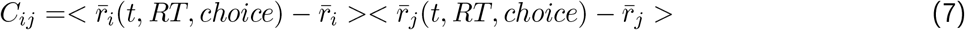

Which can be simplified (see Machens et al. (2010) for more details) to:

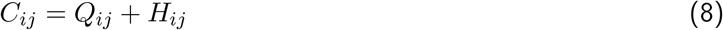

Where *Q*_*ij*_ is a signal covariance and *H*_*ij*_ is the noise covariance.

Our goal is to perform PCA on *Q*_*ij*_. However, because our data were not collected simultaneously, we cannot calculate *Q*_*ij*_ as we do not have a good estimate of *H*_*ij*_.

Nevertheless, even with trial-averaged data, one can provide an estimate of *H*_*ij*_ by constructing putative noise matrices based on the simplifying assumption that the noise is largely independent in neurons with perhaps modest noise correlations. To generate representative noise traces for our firing rates, notice that if one subtracts:

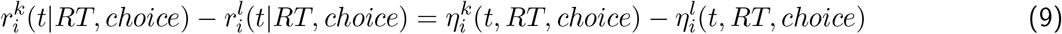

Which is just subtraction of two random instantiations of the same process, which can be written as:

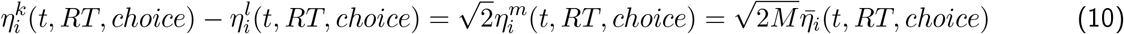

Where the final equality emerges from the equations for standard error of the mean. For example, 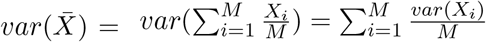.

Thus, we can generate estimates of the “average” noise 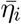(*t,RT, choice*)

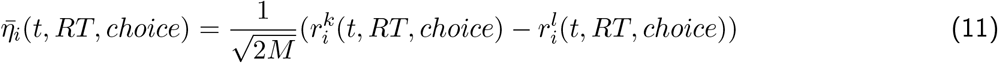

Using this equation, we can estimate *H*_*ij*_.

We denote *C*_*ij*_ as the “signal+noise” covariance matrix and *H*_*ij*_ as the “noise” covariance matrix. We estimate the eigenvalues and eigenvectors of both covariance matrices and compare them to identify the number of dimensions needed to explain the data. We used bootstrapping to derive error estimates on the signal+noise PCA and identified the number of dimensions as the first dimension where signal+noise variance was significantly below the noise variance.

### 4.11. Kinematic analysis of neural trajectories (KiNeT)

We used the recently developed KiNeT analysis (Remington et al., 2018b) to characterize how state space trajectories evolve over time in terms of relative speed and position as compared to a reference trajectory. We used the first six PCs (~90% of variance) of the PCAs organized by choice and RT/outcome as these PCs were significantly different from noise in both PCAs (Machens et al., 2010).

As such we have a collection of six-dimensional trajectories (Ω_1_, Ω_2_ … Ω_*n*_) differing in RT bins and choice in one analysis (Fig. 4C-F) and trial outcome and choice in another (Fig. 7C, E). The trajectory associated with the middle RT bin (cyan, Fig. 4C, F) and the trajectory associated with the “Correct” trial outcome (Fig. 7C, E) were chosen as ‘reference’ trajectories (Ω_*ref*_) to calculate various parameters (e.g., Time to reference) of the other non-reference trajectories (i.e., trajectories associated with the ten other RT bins and the three other trial outcomes). All of the following calculations in this section were first performed within a particular choice and then averaged across choices. Please refer to Fig. S4 for a visualization of KiNeT analyses and glossary of terms used in the following equations.

#### Time to reference

KiNeT finds the Euclidean distances between the six-dimensional position of the reference trajectory at timepoint j (*s*_*ref*_ [*j*]) and the six-dimensional position of a non-reference trajectory (Ω_*i*_) at all of its timepoints (Ω_*i*_(*τ*)). We identified the timepoint (*t*_*i*_[*j*]) at which the six-dimensional position of a non-reference trajectory (*s*_*i*_[*j*]) is closest to *s*_*ref*_ [*j*] (minimum Euclidean distance).

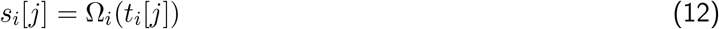

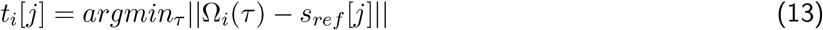

If the non-reference trajectory reaches a similar position to the reference trajectory at an ‘earlier’ timepoint then it’s a ‘faster’ trajectory (*t*_*i*_[*j*] *< t*_*ref*_ [*j*]) whereas if it reaches the same point at a ‘later’ timepoint then it is a ‘slower’ trajectory (*t*_*i*_[*j*] *> t*_*ref*_ [*j*]) (Fig. 4F).

#### Distance

The distance between reference and non-reference trajectories at timepoint j (*D*_*i*_[*j*]) is taken as the minimum Euclidean distance between the position of the reference trajectory at timepoint j (*s*_*ref*_ [*j*]) and the position of the non-reference trajectory at all its timepoints (*s*_*i*_[*j*]). Additionally, the size of the angles between a normalized non-reference trajectory and normalized trajectories for the 1st and last conditions (e.g., 1st and last RT bins) determines whether the current non-reference trajectory is closer to either the 1st or last condition. As defined here, if a trajectory is closer (i.e. smaller angle) to the trajectory for the 1st condition then (*D*_*i*_[*j*]) is positive, otherwise it is negative (Fig. 4C).

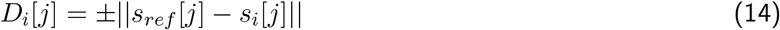

#### Angle

KiNeT computes the vector between adjacent trajectories by subtracting the positions of two non-reference trajectories when they are respectively closest to the reference trajectory at timepoint j. These vectors are then normalized and all the angles between all adjacent normalized vectors is found at all timepoints. Finally, the average angle is found at each timepoint between all adjacent trajectories (Fig. 4E).

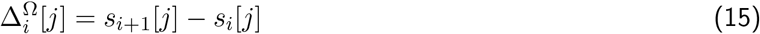

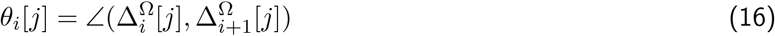

#### Subspace similarity

We first identified normalized vectors between adjacent trajectories for all timepoints. We then averaged these normalized vectors, so that we have the mean between trajectories (i.e. conditions) vector for each timepoint. This mean vector is again normalized as averaging normalized vectors doesn’t maintain unit length 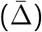. Essentially the normalized average vector is representative of the geometry of the subspace. We calculate the angle between the average vector at timepoint t (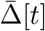 and the average vector at the first timepoint 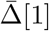, for all timepoints *t ∈ τ*. In other words we are measuring how this vector, representative of the state space, rotates relative to the first timepoint across a trial (Fig. 4D). This data is calculated separately for each choice and again is bootstrapped and averaged across the separate reaches and then across the bootstraps.

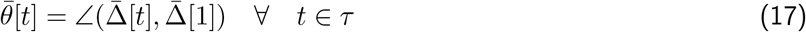

### 4.12. Scalar Speed

We computed scalar speed in firing rate state space for the prestimulus period within a RT bin (Fig. 4G) as the *𝓁*^2^ norm between the six-dimensional coordinates, of the PC data at adjacent 10 ms time steps, per RT bin and for each choice separately.

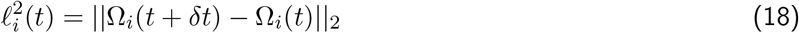

Where Ω_*i*_(*t* + *δt*) and Ω_*i*_(*t*) are six-dimensional trajectories within condition i at time t + *δt* and time t in the prestimulus period, respectively. 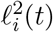 is the *𝓁*^2^ norm between six-dimensional trajectories within a condition at time t and t + 1. We then averaged speeds across choices and over the entire prestimulus period (−400 ms to 0).

The plotted ‘prestimulus firing rate speed’ was averaged across 50 bootstraps in which trials were sampled with replacement 50 times (Fig. 4G). Separate PCA and speed calculations were performed per bootstrap.

### 4.13. Choice Selectivity Signal

We estimated the ‘choice selectivity signal’ by calculating the Euclidean distance between left and right reaches at all timepoints for the first six PCs within each condition (i.e., RT bins (Fig. 4H & Fig. 6D) and coherence (Fig. 6B).

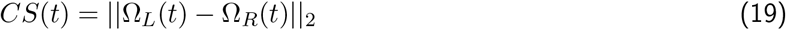

Ω_*L*_(*t*), and Ω_*R*_(*t*) - the six-dimensional location in state space for a left and right choice at time t.

To calculate the latency of this choice selectivity signal, we fit the time varying choice selectivity signal with a piecewise function of the form

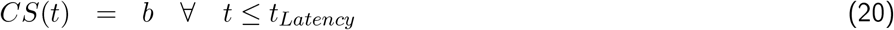

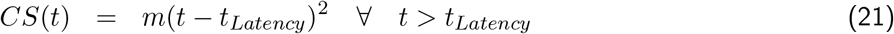

### 4.14. Initial condition as a function of RT and coherence

To estimate the initial conditions shown in Fig. 6E, F, we performed the following procedure. For each coherence and RT bin, we concatenated the average location in the six-dimensional state space in the −300 ms to −100 ms epoch before checkerboard onset for both reach directions and obtained a 77×12 matrix (7 coherences, 11 RT bins, and 2 choices). We then performed a PCA on this 77×12 matrix and used the top PC as a measure of the initial condition that we used for plotting and subsequent partial correlation analysis.

### 4.15. Latent Factors Analysis of Dynamical Systems (LFADS)

LFADS is a generative model which assumes that neuronal spiking activity is generated from an underlying dynamical system (Pandarinath et al., 2018). This dynamical system is assumed to be relatively low-dimensional (i.e. considerably smaller than the number of neurons involved) and latent factors can be extracted and exploited to recreate spiking activity on single trials. This method uses a trained autoencoder to generate ‘initial conditions’ based on a trial’s neurons’ spike counts. This ‘latent code’ serves as the initial condition to the generator RNN. From the latent code the generator infers the latent factors of all the neurons in that trial. Here LFADS was used for a single session which recorded from 23 neurons. Our model consisted of eight latent factors to recreate spiking activity of single trials. Since these factors are not orthogonal to each other, PCA was performed on these eight factors and the first three PCs were visualized in Fig. 5A, B and Fig. S3B. Please refer to Pandarinath et al. (2018) for fuller descriptions of the LFADS method.

### 4.16. Linear regression to relate RT and firing rate, and logistic regression to decode choice

We used linear and logistic regressions (decoders) to determine the variance in RT explained by spiking activity and whether spiking activity predicted choice or past outcomes, respectively. For these analyses, we leveraged the U-probe sessions where multiple neurons were recorded from at once. For monkey T, we used 24 sessions (36,690 trials) where there was a minimum of 9 neurons (one session only has 2 neurons; otherwise all other sessions had at least 9) and a maximum of 32 neurons. For monkey O, we used 27 sessions (30,831 trials) where there was a minimum of 5 neurons in a single session and a maximum of 18. Some sessions had distinct portions (e.g., the electrode was moved). In the later portion of three sessions, 2 neurons were recorded from and in another, 3 neurons were recorded from. Otherwise in all other sessions at least 5 neurons were recorded from. Variance explained and decoding accuracy shown in Fig. 5 is pooled across both monkeys.

For regression and decoding analyses, we used 1800 ms of spiking activity from each trial (600 ms prestimulus and 1200 ms poststimulus). We binned the spike times. For the choice decoder, we used 20 ms nonoverlapping bins. For the outcome decoder, we used 50 ms overlapping (10 ms time step) bins. This provided us with 90 timepoints for the choice decoder, and 72 timepoints for the outcome decoder across all units within a session.

#### Linear Regression

For analysis of the relationship between activity in PMd and RT, we regressed spike counts for each bin for all trials across all units for that session to RT according to the following equation:

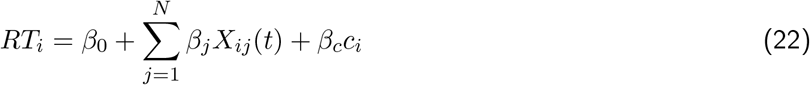

Where *RT*_*i*_(*t*) is the RT on the *i*^*th*^ trial, *X*_*ij*_(*t*) is the spike count in a 20 ms bin for the *i*^*th*^ trial and the *j*^*th*^ unit, *c*_*i*_ is the coherence for the *i*^*th*^ trial, and the *β*_*j/c*_ are coefficients for the model. After regression, we calculated variance explained by spiking activity and coherence together for each bin by using the standard equation for variance explained.

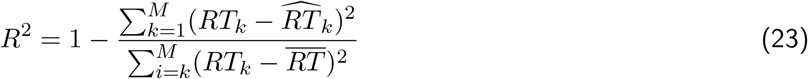

Where 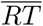 is the mean RT, *RT*_*k*_ is the RT for the *k*^*th*^ trial, and 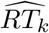 is the RT predicted for the *k*^*th*^ trial.

For assessing if the *R*^2^ values were significant, we computed a shuffled distribution (500 shuffles) where we shuffled the trials to remove the relationship between the RTs and spiking activity. We then assessed if the per bin *R*^2^ values were significantly different from the 99th percentile of the shuffled distribution *R*^2^ values.

#### Logistic Regression to decode choice

For decoding choice and previous outcome on a bin-by-bin basis, we used a regularized logistic regression approach. Decoders were trained with equal number of trials for the opposing outcomes (i.e., left vs. right reaches; previous correct vs. previous error trials). The logistic regression approach assumes that the log odds in favor of one event (e.g., left) vs. right reach is given by the following equations:

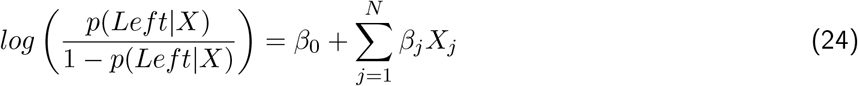

*β*_0_ - intercept of the model, *β*_*j*_ - model coefficient for the *j*_*th*_ neuron in the current bin, *X*_*j*_ - spiking activity of the *j*_*th*_ neuron of the current bin. The following equation is used to produce the outputs of the system: if *p*(*Left*|*X*) *<* 0.5 then −1 and if *p*(*Left*|*X*) *>* 0.5 then 1.

We used the implementation provided in MATLAB via the fitclinear function and the Broyden-Fletcher-Goldfarb-Shanno quasi-Newton algorithm to find the optimal fit for the parameters (Shanno, 1970). We typically attempted to predict choice or previous outcome using tens of units. To simplify the model, decrease collinearity of the coefficients and to avoid overfitting, we used L2 regularization (ridge regression):

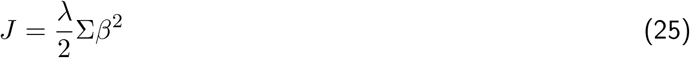

Where *J* - cost associated with coefficients, *λ* - penalty term (1/number of in-fold observations), and *β* are the coefficients of the model. We used 5-fold cross validation and calculated loss for each model. Accuracy is reported as *accuracy* = 1 − *mean*(*loss*).

### 4.17. Subspace overlap analysis

To determine how much of firing rate covariance with RTs could be explained by the outcome subspace (PCA organized by choice and outcome, Fig. 7A, B) we performed an analysis where firing data from all 996 units organized by RT and choice was projected into the first six dimensions from the PCA organized by outcome and choice. For this purpose we used a modified version of the alignment index developed by Elsayed et al. (2016):

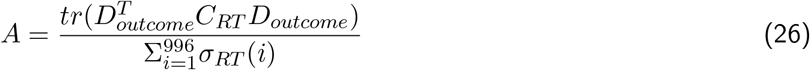

The alignment index, A, provides an estimate of the fraction of variance that is explained by projecting one subspace into another. *tr*() is the trace of a matrix, which can be proved to be the sum of its eigenvalues. *D*_*outcome*_ is the first six eigenvectors of all 996 units from the PCA organized by outcome and choice. *C*_*RT*_ is the covariance matrix of the firing rates of all 996 units organized by RT and choice. *σ*_*RT*_ (*i*) are the eigenvalues (i) of the covariance matrix organized by RT and choice. For our purposes, we used the total variance in the denominator instead of the same number of dimensions as the numerator. Thus, the alignment index calculates the ratio of how much of the *total variance* from firing data organized by RT and choice is explained by the outcome subspace.

### 4.18. Demixed principal component analysis (dPCA)

We used dPCA, a semi-supervised dimensionality reduction technique to further understand if prestimulus activity which covaried with RTs shared variance with firing rate activity that covaried with the previous trial’s outcome. We performed two dPCAs. The first identified axes that maximally accounted for firing rate variability from trial outcome and the second identified axes that maximally accounted for firing rate variability that covaried with RTs. We then calculated the dot product between these axes and estimated the angle using the inverse cosine of the dot product. An angle of zero would indicate that these axes completely overlap and that their sources of variance are the same, whereas orthogonal angles would mean that the axes do not overlap and therefore share no variance.

## 5. Author Contributions

CC trained both monkeys and recorded in PMd using multi-contact electrodes under the mentorship of KVS. PB, TW, and CC jointly collaborated on the various analyses. GK and LC provided helpful insights for analysis and relevant literature for the manuscript. PB and CC wrote initial drafts of the paper. All authors refined further drafts contributing analyses, insights, and writing.

## 6. Acknowledgments

We thank Dr. Michael Economo, Dr. Kamal Sen, and Dr. Matt Golub for comments on the previous versions of the manuscript.

CC was supported by a NIH/NINDS R00 award R00NS092972, the Moorman-Simon Interdisciplinary Career Development Professorship from Boston University, the Whitehall foundation, the Young Investigator Award from the Brain and Behavior Research Foundation, and an NIH/NINDS R01 award NS122969. KVS was supported by the following awards: NIH Director’s Pioneer Award 8DP1HD075623, NIDCD R01-DC014034, NIDCD U01-DC017844, NINDS UH2-NS095548, NINDS UO1-NS098968, DARPA-BTO ‘REPAIR’ Award N66001-10-C-2010, DARPA-BTO ‘NeuroFAST’ award W911NF-14-2-0013, Simons Foundation Collaboration on the Global Brain awards 325380 and 543045, Office of Naval Research award N000141812158, Larry and Pamela Garlick, Wu Tsai Neurosciences Institute at Stanford, the Hong Seh and Vivian W. M. Lim endowed professorship and the Howard Hughes Medical Institute. The funders had no role in study design, data collection and interpretation, or the decision to submit the work for publication.

## Supplemental materials

**Figure S1:**
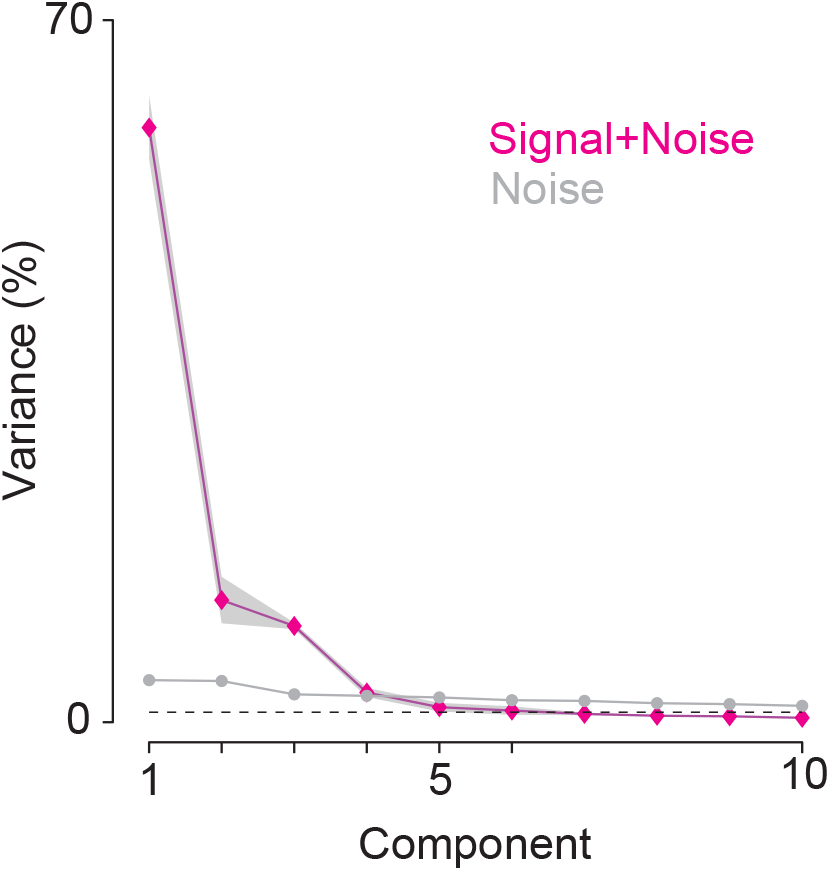
Percent variance explained by each component from the PCA organized by RT and choice: “signal+noise” and “noise” variance explained by the first 10 components. The first six components capture over 90% of the variance. To derive the error bars for the signal+noise PCA, we used bootstrapping (50 repeats) over trials to estimate standard errors.

**Figure S2:**
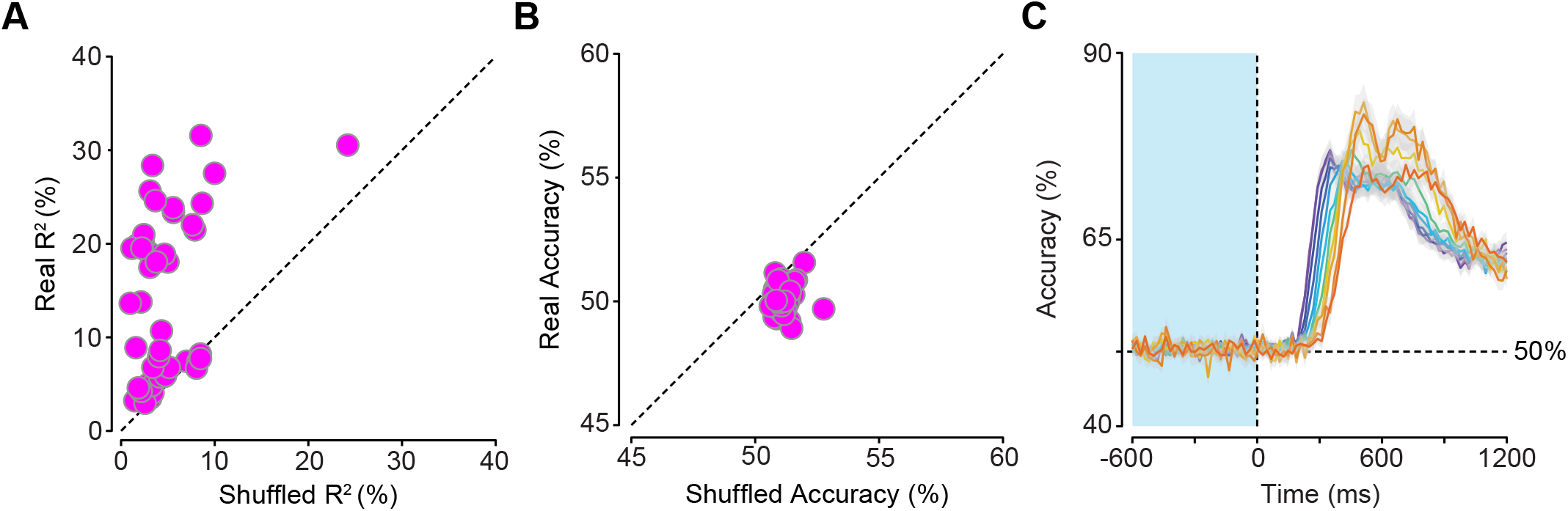
Prestimulus spiking activity is predictive of RT but not choice, even when decoding is performed within RT bins. (**A/B**) Scatterplot of true mean prestimulus *R*^2^/accuracy values compared to the *R*^2^/accuracy values for the 99th percentile of the shuffled data. Each dot represents the bin-and trial-averaged prestimulus mean *R*^2^/accuracy value within each of the 51 sessions. The dotted line is where scatter points would fall if shuffled *R*^2^ and real *R*^2^ values were equivalent. (**A**) Many of the points lie above this line suggesting that real prestimulus neural activity explains more of the RT variance than shuffled neural data. (**B**) In contrast, many of the points lie on or below this line suggesting that real prestimulus neural activity is not predictive of choice. (**C**) Plot of mean accuracy from logistic regressions of binned spiking activity (20 ms) used to predict trial-matched eventual choice within RT bins. Accuracy is averaged across 51 sessions. Gray shaded area is *SEM*. The gray dotted line is 50% accuracy.

**Figure S3:**
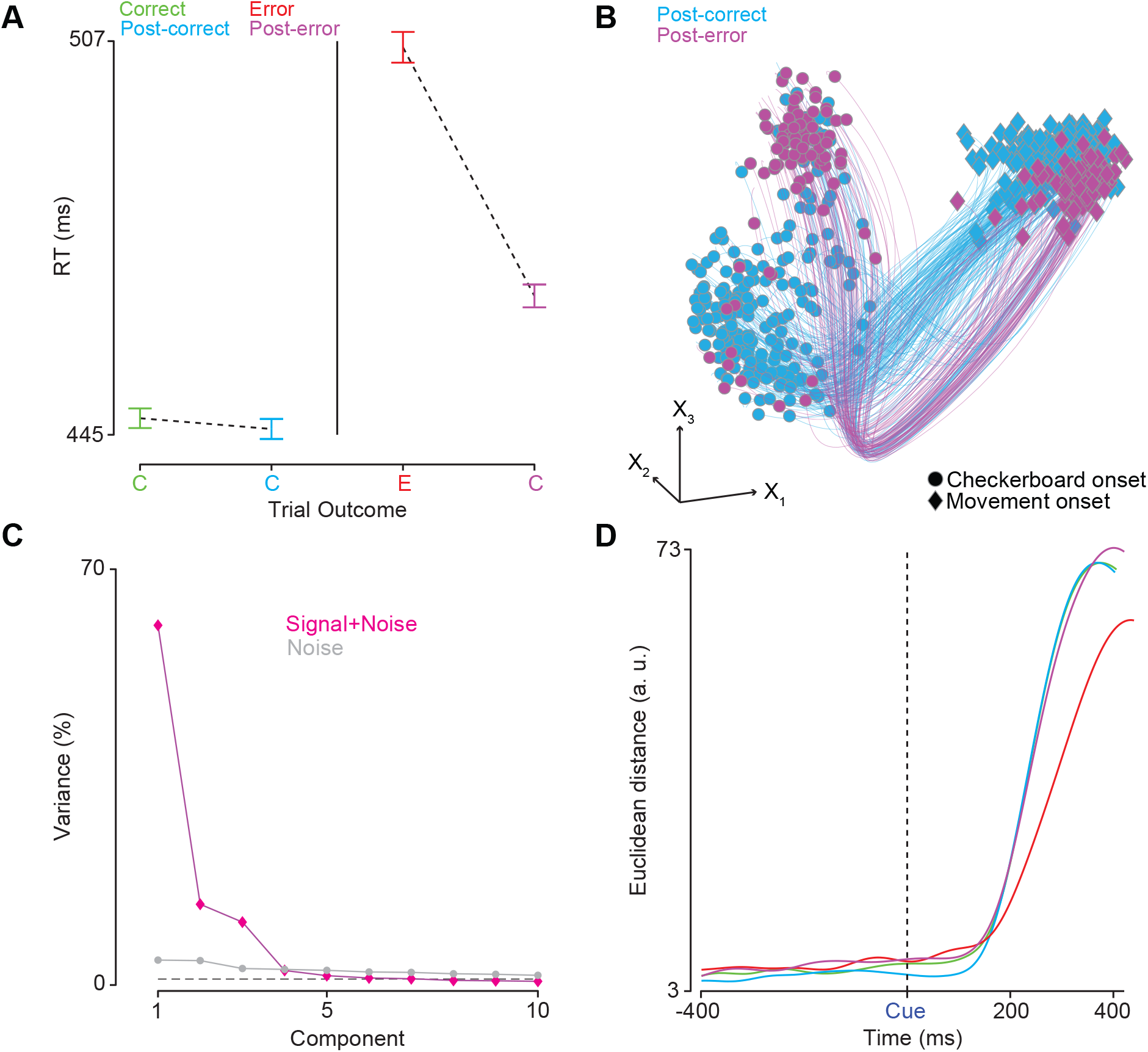
Neural dynamics associated with post-error slowing may demonstrate larger choice selectivity. (**A**) Average RTs from all error → correct and trial-matched correct → correct sequences found across both monkeys and all sessions (error bars are 2 × SEM). On average both monkeys demonstrate classical post-error slowing. (**B**) LFADS trajectories in the space of the first three orthogonalized factors (*X*_1,2,3_), obtained via PCA on LFADS latents, for 30% of post-correct and all post-error trials, for all coherences and left reaches from a single session (23 units). Each trajectory is plotted from 200 ms before checkerboard onset (dots) to movement onset (diamonds). (**C**) Scree plot of the percentage of variance explained by the first ten components. The first six PCs capture ~90% of the variance in firing rate activity. (**D**) Euclidean distance in the first six dimensions between the two reach directions aligned to checkerboard onset (‘Cue’ & black dashed line). We observed no prestimulus separation between reach directions. Choice selectivity is lower and slower for error trials compared to all other outcomes. Post-error choice selectivity may be larger than other trial outcomes.

**Figure S4:**
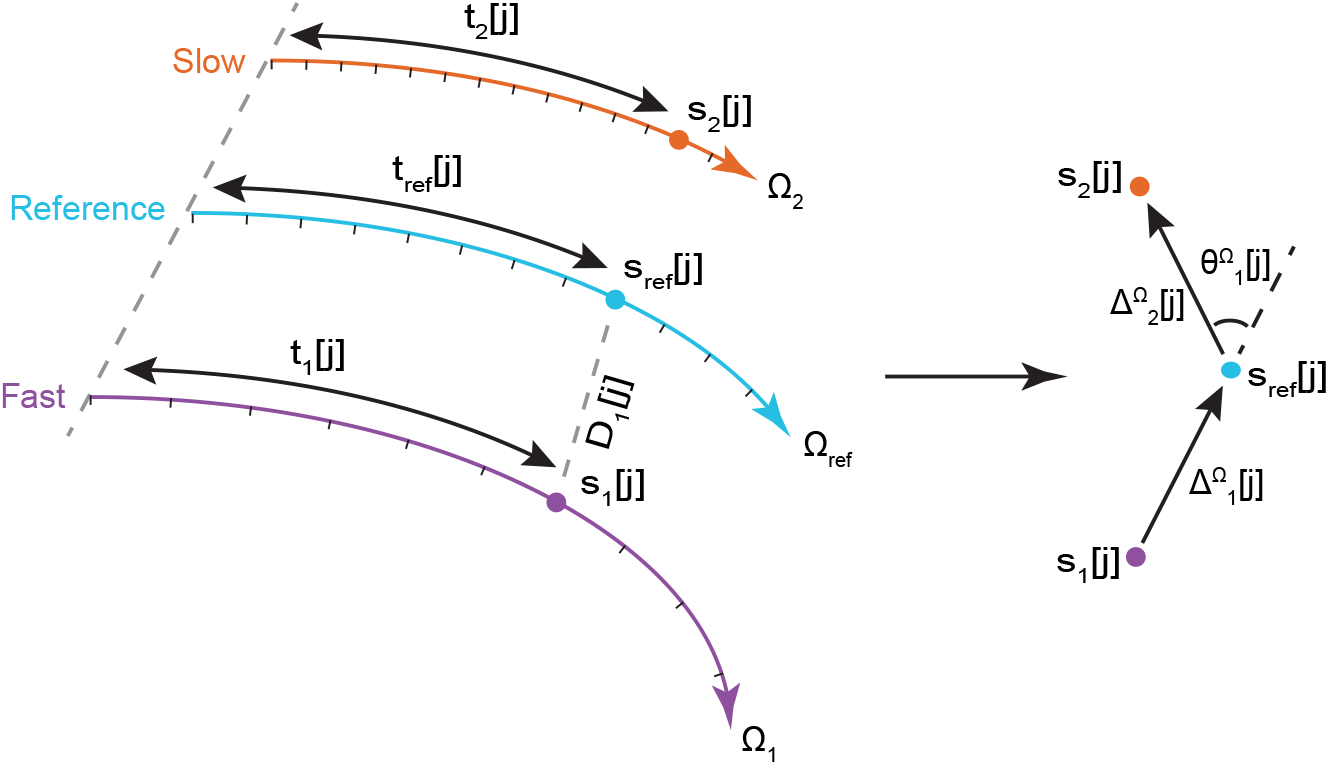
Diagram of kinematic analysis of neural trajectories (KiNeT) The middle trajectory in cyan denotes the reference trajectory Ω_*ref*_. Two non-reference Ω_1_ (violet) and Ω_2_ (orange) denote trajectories that evolve faster and slower than Ω_*ref*_, respectively. For each timepoint j, the corresponding neural state on the reference trajectory is denoted as *s*_*ref*_ [*j*] and *t*_*ref*_ [*j*] is the corresponding time for the reference trajectory to evolve from the initial point to *s*_*ref*_ [*j*]. The closest points to *s*_*ref*_ [*j*] on the fast and slow trajectories as measured by Euclidean distance are denoted by *s*_1_[*j*] and *s*_2_[*j*]. *t*_1_[*j*] and *t*_2_[*j*] are the corresponding times to reach *s*_1_[*j*] and *s*_2_[*j*]. The vector connecting the two closest points on adjacent trajectories at timepoint j is denoted as Δ*i*^Ω^[*j*]. The angle between two adjacent vectors is calculated as *θ*_*i*_[*j*] • *i* - index of non-reference trajectories • *j* - index of timepoints associated with the reference trajectory • Ω_*i*_ - the *i*^*th*^ non-reference trajectory • *τ* - timepoints associated with the non-reference trajectories • Ω_*i*_(*τ*) - position of non-reference trajectory at timepoint *τ* • *s*_*ref*_ [*j*] - position of reference trajectory at the *j*^*th*^ timepoint • *s*_*i*_[*j*] - closest position of Ω_*i*_ to *s*_*ref*_ [*j*] at timepoint j • *t*_*i*_[*j*] - the non-reference timepoint for when it’s closest to the reference trajectory at timepoint j (corresponding time of *s*_*i*_[*j*]) • *D*_*i*_[*j*] - distance between nearest point on non-reference trajectory Ω_*i*_ and reference trajecory at index j • Δ*i*^Ω^[*j*] - vector connecting two nearest points on two adjacent trajectories. • *θ*_*i*_[*j*] - angle between two adjacent vectors 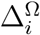 and 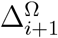 • *argmin* - where function achieves its minimum at point *j*

